# Natural variation in transplacental transfer efficiency exposes distinct transcriptional network architectures of PFAS effects on birth weight and gestational age

**DOI:** 10.64898/2026.03.23.712893

**Authors:** Sean T. Bresnahan, Hannah E. J. Yong, Maggie G. Drelichman, Soraya N. Campbell, Ance E. Trapse, Gabriel R. Romo, Christopher M. Cellini, Sierra Lopez, Jerry Kok Yen Chan, Shiao-Yng Chan, Elana R. Elkin, Arjun Bhattacharya, Jonathan Y. Huang

**Affiliations:** Department of Epidemiology, University of Texas MD Anderson Cancer Center, Houston, TX, USA; A*STAR Institute for Human Development and Potential, Singapore, Singapore; Division of Environmental Health, School of Public Health, San Diego State University, San Diego, CA, USA; Department of Public Health Sciences, The University of Hawai’i at Mānoa, Honolulu, HI, USA; KK Women’s and Children’s Hospital, Singapore, Singapore; Duke-NUS Medical School, Singapore, Singapore; Department of Obstetrics and Gynecology, Yong Loo Lin School of Medicine, National University of Singapore, Singapore, Singapore; Institute for Data Science in Oncology, University of Texas MD Anderson Cancer Center, Houston, TX, USA

## Abstract

Per- and polyfluoroalkyl substances (PFAS) are environmental contaminants that produce heterogeneous effects on perinatal outcomes despite chemical similarity. Natural variation in transplacental transfer efficiency (TPTE, the degree to which compounds cross the placental barrier) presents a mechanistic lens for understanding this heterogeneity, but whether TPTE systematically shapes the transcriptional pathways linking exposure to outcomes has not been tested. Using isoform-resolved placental transcriptomics from n = 124 term deliveries with eight PFAS measured in maternal and cord blood, validated against patient-derived placental explants (n = 18) using a tissue-specific long-read transcriptome assembly, we show that PFAS influence perinatal outcomes primarily through co-expression network hubs rather than differentially expressed features, indicating that network position, not fold-change magnitude, determines mediator importance. Leveraging TPTE as a natural experiment, we find that direct fetal exposure recruits increasingly numerous and tightly coordinated transcriptional mediators for both birth weight and gestational age, but the two outcomes diverge in network architecture: network centrality and maternal-fetal compartmentalization scale with TPTE exclusively for birth weight, while gestational age shows no such topological reorganization. These outcome-specific patterns are detectable only at the transcript level, as gene-level aggregation masks systematic TPTE-network relationships. This framework distinguishes which perinatal outcomes are mechanistically vulnerable to fetal exposure dose, with implications for risk assessment and therapeutic target identification for environmental contaminants.

## Introduction

Per- and polyfluoroalkyl substances (PFAS) are a class of synthetic organofluorine compounds characterized by strong carbon-fluorine bonds that confer exceptional chemical stability and resistance to environmental degradation.^1^ These properties have made PFAS ubiquitous in commercial and industrial applications, including food packaging, non-stick cookware, water-repellent textiles, and firefighting foams.^2^ However, their persistence and bioaccumulative potential have resulted in widespread environmental contamination and nearly universal human exposure.^3^ Biomonitoring studies consistently detect PFAS in the serum of over 95% of the U.S. population, with concentrations varying by geographic region, dietary habits, and occupational exposures.^4,5^

The critical need to study PFAS stems from mounting evidence of their adverse health effects across multiple organ systems and life stages. PFAS exposure is associated with immunotoxicity, metabolic dysfunction, hepatotoxicity, endocrine disruption, and increased cancer risk.^1,6^ Of particular concern is the vulnerability of developing fetuses to PFAS exposure, as these compounds readily cross the placental barrier.^7^ The capacity of PFAS to accumulate in maternal tissues and transfer to the fetus during critical windows of development raises substantial concerns about long-term health consequences, including potential transgenerational effects.^8–10^

During gestation, PFAS exposure has been linked to multiple adverse pregnancy outcomes and developmental effects. Epidemiological studies have consistently reported associations between maternal PFAS concentrations and reduced birth weight, with effect sizes varying by specific PFAS compound.^11^ Meta-analyses indicate that prenatal exposure to perfluorooctanoic acid (PFOA) and perfluorooctane sulfonate (PFOS), two of the most studied PFAS, are associated with decreases in birth weight ranging from 10-20 grams per natural log unit increase in maternal serum concentration.^12^ Beyond fetal growth restriction, PFAS exposure during pregnancy has been implicated in preeclampsia, gestational diabetes, preterm birth, and altered immune function in offspring.^13,14^

Notably, birth weight and gestational age at delivery, two key perinatal outcomes affected by PFAS, represent mechanistically distinct endpoints. Birth weight reflects fetal growth and size, primarily determined by placental nutrient transfer, fetal cellular proliferation, and growth factor signaling. In contrast, gestational age reflects the timing of parturition, governed by maternal-fetal endocrine signaling, prostaglandin synthesis, inflammatory cytokine cascades, and myometrial contractility.^15^ Critically, small-for-gestational-age (birthweight < 10^th^ percentile for gestational age) can occur at any gestational age, and preterm birth (< 37 weeks’ gestation) can occur with low, appropriate or excessive fetal growth, demonstrating the independence of these regulatory systems. Despite a general correlation between birthweight and gestational age, few studies have convincingly demonstrated differential mechanisms for PFAS effects on these phenotypes, and whether environmental toxicants differentially perturb pathways controlling fetal growth versus parturition timing remains incompletely understood.

The placenta serves as the critical interface between maternal and fetal circulations, mediating nutrient transport, waste exchange, hormone production, and immune protection throughout pregnancy.^16^ As such, the placenta represents a unique organ for investigating the mechanisms by which environmental toxicants affect fetal development. Studies examining PFAS in placental tissue have revealed compound-specific patterns of accumulation and differential transplacental transfer efficiencies (TPTE), defined as the ratio of fetal to maternal circulating concentrations.^7,17,18^ Shorter-chain PFAS such as perfluorobutane sulfonate (PFBS) exhibit higher TPTE values compared to longer-chain compounds like PFOS, suggesting that placental handling varies by molecular structure.^19^ Variation in TPTE across compounds provides a natural experiment for investigating how the degree of direct fetal exposure modulates placental transcriptional responses and whether compounds achieving greater fetal exposure engage distinct regulatory programs controlling different perinatal outcomes.

At the molecular level, limited transcriptomic studies have identified altered expression of genes involved in lipid metabolism, oxidative stress responses, and hormone signaling in placentas from pregnancies with elevated PFAS exposure.^20^ However, these studies have been constrained by reliance on gene-level rather than isoform-level resolution, which fails to capture alternative splicing and isoform usage that may be particularly susceptible to toxicant exposures.^21^ Previous work has demonstrated that isoform-level modeling increases discovery of transcriptome-mediated associations with complex traits by approximately 60% compared to gene-level analyses, as alternative splicing patterns and isoform usage explain substantially more phenotypic variance than aggregated gene expression.^22–25^

Beyond resolution, existing studies have primarily focused on identifying differentially expressed features without determining whether they causally mediate environmental effects on perinatal outcomes or how they are organized within broader regulatory networks. Standard differential expression analyses implicitly assume that fold-change magnitude reflects functional importance, yet genes under the strongest dosage constraint, e.g., transcription factors and developmental regulators, tolerate the least expression variation, making the most functionally critical genes often show the smallest detectable changes.^25,26^ Empirical studies demonstrate that network hub genes are approximately three times more likely to be essential than peripheral genes,^27,28^ and in developmental contexts, hubs with positive feedback loops amplify weak perturbations into switch-like triggers that influence cell fate decisions.^29^ This regulatory leverage is particularly relevant for environmental toxicants like PFAS that produce subtle but coordinated transcriptomic perturbations through nuclear receptor or transcription factor pathways.^30,31^ In placenta studies specifically, weighted co-expression network analysis (WGCNA)-identified hub genes have been validated as statistical mediators of exposure–birth outcome relationships even when they fail significance thresholds in standard differential expression analysis,^32,33^ with co-expression modules shown to mediate 31–61% of the relationship between prenatal exposure to another group of chemicals, phthalates, and placental efficiency measures.^34^ Understanding PFAS toxicity mechanisms therefore requires distinguishing between transcripts showing the largest expression changes and those occupying central positions within co-expression networks that coordinate physiological responses and lie on causal pathways between exposure and adverse outcomes.

In this study, we leverage isoform-resolved placental transcriptomics from the Growing Up in Singapore Towards healthy Outcomes (GUSTO) prebirth study with measured concentrations of eight PFAS compounds in both maternal delivery blood and cord blood^17,35^ to differentiate mechanisms underlying PFAS effects on birth weight and gestational age. Using a tissue-matched long-read transcriptome reference^36^, we demonstrate improved concordance between epidemiological findings and experimental validation in patient-derived placental explants, with PFAS-associated differential expression manifesting more extensively at the isoform level. Through integration of causal mediation analysis and weighted co-expression network analysis, we find that PFAS effects on perinatal outcomes are mediated primarily through network hubs rather than differentially expressed features, with the strength, network topology, and compartmentalization of mediation effects scaling systematically with TPTE in outcome-specific patterns **(Figure 1)**. These findings reveal how natural variation in TPTE resolves mechanistically distinct placental responses underlying PFAS effects on fetal growth versus parturition timing, with implications for PFAS exposure-associated disease etiology, risk assessment, and identification of therapeutic targets.

**Figure 1.**
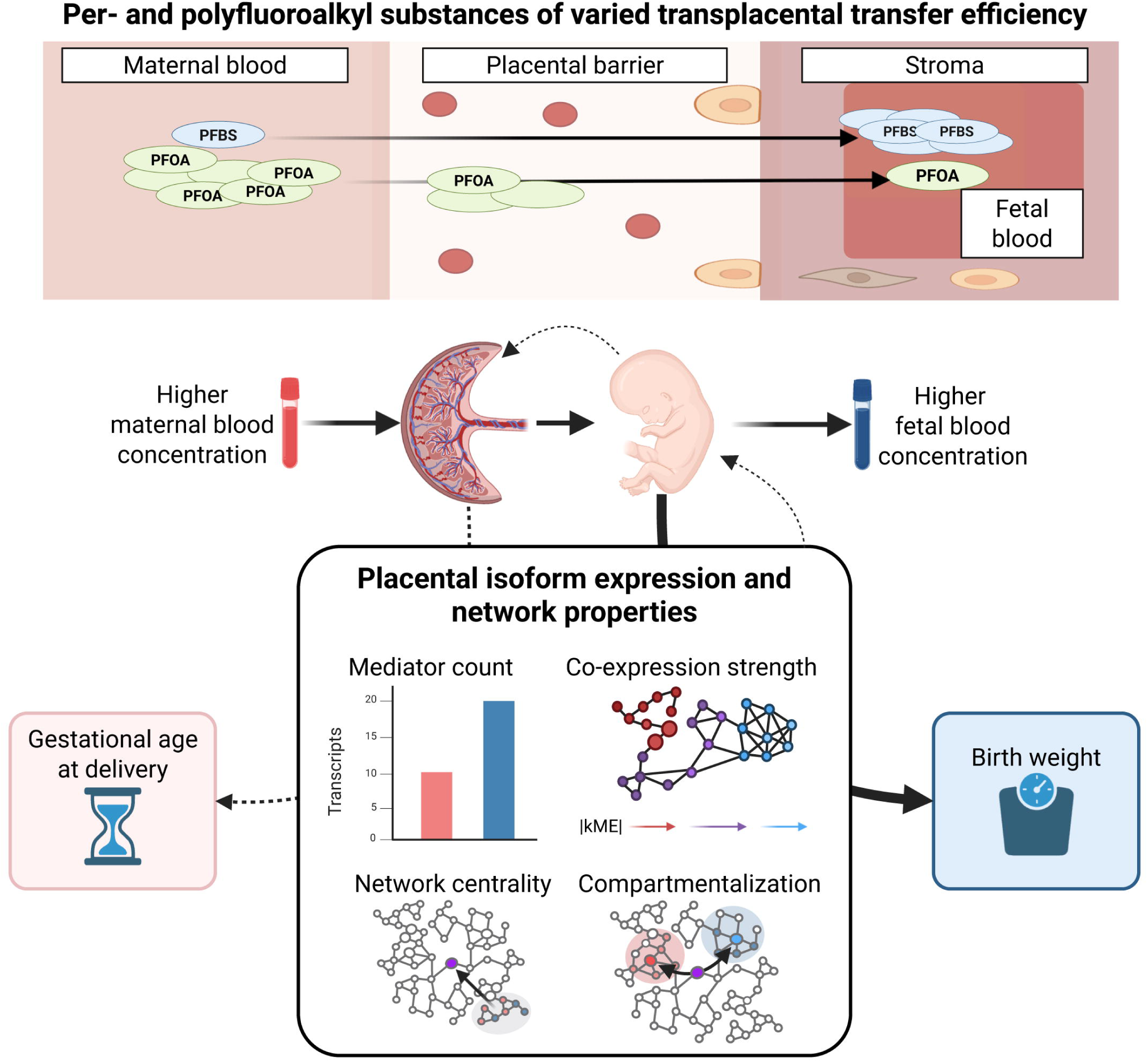
Transplacental transfer efficiency reveals dose-dependent network architectures linking PFAS exposure to birth outcomes. Per- and polyfluoroalkyl substances (PFAS) exhibit varied transplacental transfer efficiencies (TPTE), quantified as the ratio of fetal cord blood to maternal blood concentrations. PFOS shows low TPTE with limited fetal exposure, while PFBS demonstrates high TPTE with substantial transfer to the fetus. PFAS concentrations were measured in maternal blood and fetal cord blood alongside placental transcriptomics, gestational age at delivery, and birth weight. Placental transcripts mediating PFAS effects on birth weight were characterized by four network properties: mediator count (number of significant mediating transcripts), co-expression strength (|kME|, representing correlation with module eigengene and shown as network connectivity), network centrality (hub positioning within the network), and compartmentalization (spatial clustering of maternal versus fetal mediators). Birth weight exhibited coordinated TPTE-dependent responses across these network metrics, with high-TPTE compounds engaging more numerous, strongly co-expressed mediators occupying central network positions with distinct maternal-fetal compartmentalization. In contrast, gestational age showed minimal coordinated network reorganization in response to TPTE variation, indicating distinct molecular architectures underlying PFAS effects on these different outcomes.

## Results

### Long-read transcriptome assembly enhances concordance between observational and experimental PFAS effects on placental transcript expression

To investigate the molecular mechanisms through which PFAS exposure affects birth weight, we analyzed fetal-facing placental tissue samples from term deliveries (n = 124) in the GUSTO prebirth study, a cohort of predominantly East, Southeast, and South Asian ancestry, with measured concentrations of eight PFAS compounds in maternal delivery blood and/or cord blood^17,35^ **(Table 1**; **Figure 2a)**. We quantified short-read RNA-seq data from GUSTO samples against both isoform-resolved placental transcriptome reference we previously assembled from long-read and orthogonal sequencing data^36^ and the GENCODEv45 transcriptome reference. To assess whether reference annotation choice modifies effect estimates in observational studies, we compared differential expression patterns from the GUSTO cohort with those from an independent cohort of patient-derived placental explants sampled at two developmental windows (second trimester and term) and treated with PFBS at 0 µM, 5 µM, and 20 µM concentrations (n = 18; n = 3 per group), modeling PFBS concentration as a continuous exposure for 24 hours.

**Figure 2.**
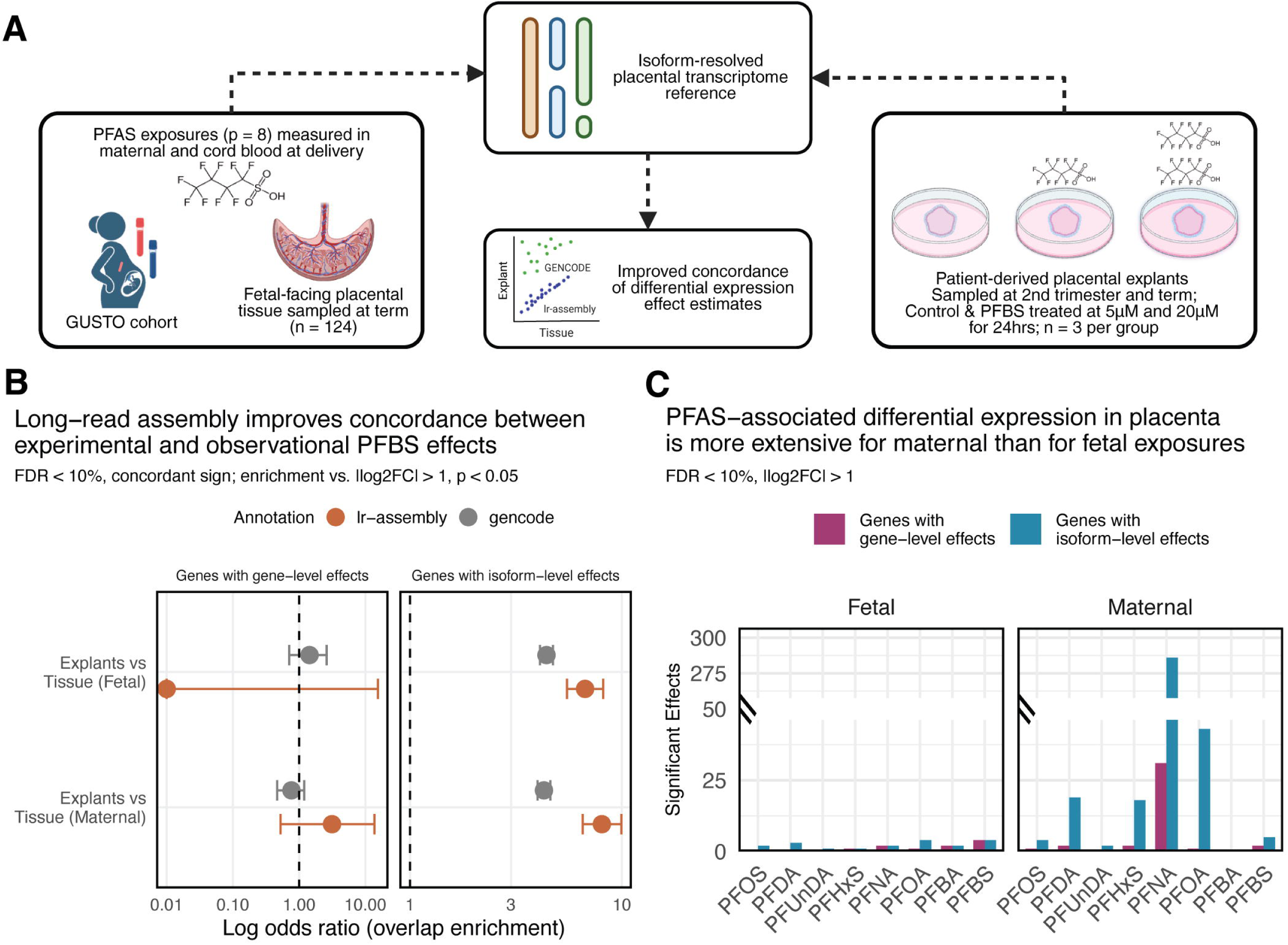
Long-read based transcriptome assembly of placenta enhances concordance between experimental and observational PFAS effect estimates with short-read sequencing data. **(A)** Conceptual diagram of the study design integrating observational prebirth study data with experimental placental explant models to investigate PFAS-responsive transcriptional changes. Fetal-facing placental tissue (GUSTO, n = 124; n = 101 with fetal PFBS, n = 101 with maternal PFBS, n = 78 with both) from the GUSTO prebirth study cohort with PFAS exposures (p = 8 compounds) measured in maternal blood collected at delivery (pink vial) and cord blood (blue vial) was analyzed using an isoform-resolved placental transcriptome reference. PFBS-associated differential expression patterns observed in the GUSTO samples were validated using patient-derived placental explants sampled at second trimester and term and treated with PFBS at 5 µM and 20 µM concentrations for 24 hours (n = 18; n = 3 per group), demonstrating improved correspondence of transcriptional responses between experimental and observational studies. **(B)** Log odds ratios for overlap enrichment between genes showing concordant differential expression with PFBS exposure in explant models compared to term placental tissue, based on long-read assembly (terracotta) or GENCODE (grey) reference annotations. The concordant overlap set was defined as nominally significant (p < 0.05) features with the same direction of effect in both datasets, and enrichment was tested against features nominally significant in explants (|log_2_FC| > 1, p < 0.05) using Fisher’s exact test. For isoform-level effects, overlap is assessed at the gene level: genes are counted if they harbor at least one significantly differentially expressed transcript in both datasets. Results are shown separately for fetal (cord blood) and maternal blood PFBS exposures. Models for placental tissue adjust for fetal sex and gestational age, while models for placental explants adjust for trimester (explants derived from 2^nd^ trimester and term samples). Both models adjust for technical variance estimated with RUVr. **(C)** Number of genes showing significant differential expression (|log_2_FC| > 1, FDR < 10%) in term placental tissue samples at gene-level (plum) or isoform-level (tefal) across eight PFAS compounds, separated by fetal (cord) and maternal blood exposures. All models adjust for fetal sex, gestational age and technical variance.

**Table 1.** Characteristics of GUSTO prebirth study participants with placental RNA-seq and PFAS measurements, stratified by labor onset.

|  | Overall (n = 124) | Spontaneous,<br>regardless of<br>delivery mode<br>(n = 72) | Other/Unknown†<br>(n = 52) |
| --- | --- | --- | --- |
| ***Fetal characteristics*** |  |  |  |
| Fetal sex, male, n (%) | 60 (48.4) | 37 (51.4) | 23 (44.2) |
| Gestational age (weeks),<br>median [IQR] | 39.0 [38.0–39.7] | 39.1 [38.0–39.7] | 38.9 [38.0–40.0] |
| Birth weight (g), median [IQR] | 3029 [2821–3350] | 3022 [2829–3285] | 3055 [2778–3452] |
| ***Maternal characteristics*** |  |  |  |
| Age at delivery (years),<br>median [IQR] | 31.9 [28.2–35.9] | 32.3 [28.5–36.0] | 30.9 [27.7–35.8] |
| ***Maternal ethnicity, n (%)*** |  |  |  |
| Chinese | 79 (63.7) | 45 (62.5) | 34 (65.4) |
| Malay | 23 (18.5) | 16 (22.2) | 7 (13.5) |
| Indian | 22 (17.7) | 11 (15.3) | 11 (21.2) |
| ***PFAS measurement availability, n (%)*** |  |  |  |
| Cord blood (fetal) only | 23 (18.5) | 14 (19.4) | 9 (17.3) |
| Maternal blood only | 23 (18.5) | 16 (22.2) | 7 (13.5) |
| Both cord and maternal blood | 78 (62.9) | 42 (58.3) | 36 (69.2) |

*IQR, interquartile range; PFAS, per- and polyfluoroalkyl substances.*
† Other/Unknown includes induced labor (n = 35) and deliveries with unrecorded onset (n = 17).
PFAS were measured in cord blood (fetal) and/or maternal blood at delivery. Of 124 participants, 78 (62.9%) had paired maternal and cord blood measures.

Long-read transcriptome assembly substantially improved concordance between observational and experimental findings for isoform-level effects. When examining features showing concordant differential expression with PFBS exposure in both datasets (nominal p < 0.05, same direction of effect), enrichment among features nominally significant in explants (|log_2_FC| > 1, p < 0.05) was similar for gene-level effects regardless of reference annotation, but markedly improved at the isoform level when using the long-read assembly (6.7- to 8.0-fold enrichment between exposure sources) versus GENCODE (4.3- to 4.4-fold enrichment) annotations **(Figure 2b; Supplementary Figure 1a)**. The strength of correlation between explant and tissue effect sizes increased with more stringent thresholds in the explant models, with R^2^ values plateauing at |log_2_FC| > 1.0 for both analyses **(Supplementary Figure 1b)**. This pattern was consistent across both fetal and maternal PFBS exposures, underscoring the value of isoform resolution in capturing environmentally responsive transcriptional changes.

Isoform-level analysis revealed more extensive differential expression than gene-level analysis across all eight PFAS compounds. In term placental tissue, isoform-level analysis detected 2- to 5-fold more differentially expressed transcripts compared to gene-level aggregation (|log_2_FC| > 1, FDR < 10%), with this pattern holding for both maternal and fetal exposure sources **(Figure 2c)**. Differential expression was consistently more extensive for maternal PFAS exposures compared to fetal exposures across compounds, suggesting that maternal blood concentrations quantified at the time of delivery may capture a broader window of placental exposure or that maternal physiological responses to PFAS have more pronounced transcriptional effects on placental function. Together, these findings suggest that isoform-resolved transcriptomics, validated through experimental concordance, provides enhanced sensitivity for detecting PFAS-associated molecular changes in the placenta.

### PFAS compounds show outcome-specific transcriptional mediation of birth weight and gestational age despite phenotypic correlation

The distinct differential expression patterns between maternal and fetal exposure sources prompted us to investigate associations between PFAS exposures, placental transcription, and perinatal outcomes **(Figure 3a)**. Birth weight and gestational age at delivery are positively correlated (r = 0.426, R^2^ = 0.182, p = 7.852 × 10^-7^) **(Supplementary Figure 2a)** yet represent mechanistically distinct outcomes. To identify transcriptional mediators of PFAS effects on these outcomes, we performed causal mediation analysis testing all annotated transcripts (n = 31,007) for each PFAS compound and exposure source, adjusting for fetal sex and gestational age (for birth weight analyses) or fetal sex alone (for gestational age analyses). All gestational age analyses were restricted to births following spontaneous onset of labor (regardless of mode of delivery; **Table 1**).

**Figure 3.**
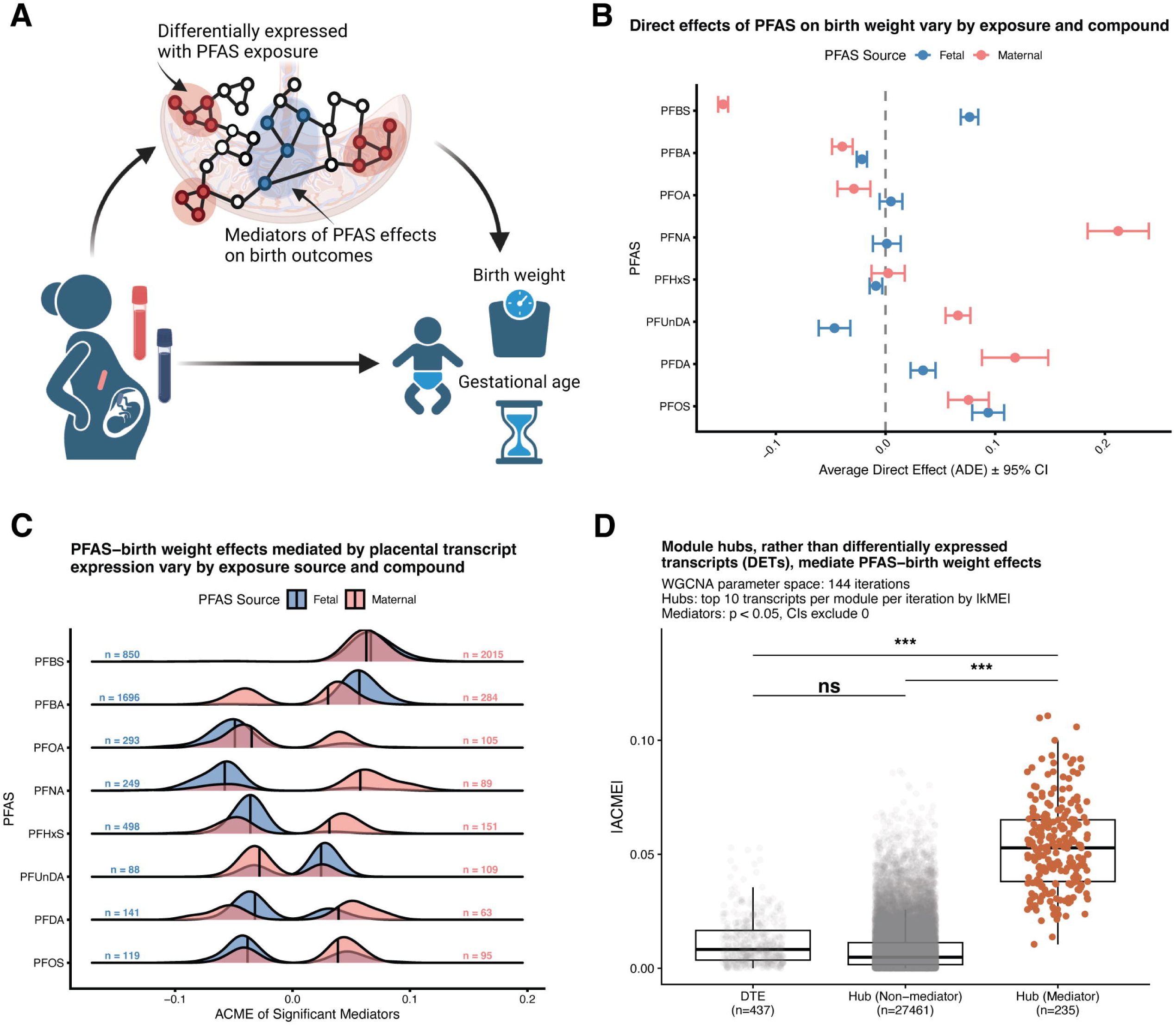
PFAS effects on birth weight are mediated through co-expression network hubs rather than differentially expressed features. **(A)** Conceptual framework for mediation analysis of PFAS effects on perinatal outcomes. Maternal PFAS exposure (measured in blood collected at delivery, pink vial) and fetal PFAS exposure (measured in cord blood, blue vial) influence perinatal outcomes including birth weight and gestational age. We hypothesized that PFAS effects operate through placental transcriptional responses, with mediation effects potentially concentrated in either differentially expressed transcripts (red nodes) or co-expression network hubs (blue nodes) that coordinate broader transcriptional programs. Mediation analysis tests whether placental transcript abundance statistically accounts for the relationship between PFAS exposure and perinatal outcomes, distinguishing direct effects of PFAS from indirect effects operating through transcriptional changes. **(B)** Average direct effects (ADE) of PFAS on birth weight vary by compound and exposure source. Mean ADE ± 95% confidence intervals for maternal blood (pink points) and fetal (cord blood, blue points) PFAS exposure sources, calculated across all significant mediation pathways. Points represent weighted mean ADEs using inverse variance weighting across significant transcript mediators. **(C)** PFAS–birth weight effects mediated by placental transcript expression vary by exposure source and compound. Density distributions of absolute average causal mediation effects (|ACME|) for significant mediators (p < 0.05 and 95% CI excludes zero) by PFAS compound and exposure source. Blue distributions represent fetal (cord blood) PFAS exposures; pink distributions represent maternal blood PFAS exposures. Vertical lines within each distribution indicate the median |ACME|. Sample sizes (n = fetal, n = maternal) indicate the number of significant mediating transcripts for each PFAS-source combination. **(D)** Module hubs rather than differentially expressed transcripts (DETs) mediate PFAS-birth weight effects. Absolute average causal mediation effects (|ACME|) for DETs (FDR < 0.1, |log_2_FC| > 1), and hubs (top 10 transcripts per module per iteration by |kME|), stratified by non-mediators and mediators (significant mediation p < 0.05, 95% CI excludes 0). Statistical comparisons via Tukey HSD following ANOVA with bootstrapping: ns = not significant, ***p < 0.001.

The average direct effect (ADE) of PFAS on birth weight varied substantially by compound and exposure source, with effects showing distinct patterns between maternal and fetal exposures **(Figure 3b)**. Similar patterns were observed for gestational age **(Supplementary Figure 2b)**. Maternal PFAS exposures generally exhibited larger direct effects compared to fetal exposures, consistent with the more extensive differential expression observed for maternal sources. We identified significant mediators (p < 0.05, 95% CI excludes zero) across all PFAS-exposure combinations, with absolute average causal mediation effects (|ACME|) varying by compound and exposure source (**Figure 3c**; **Supplementary Figure 2c** for gene-level analysis; **Supplementary Figure 2d** for gestational age). Despite the correlation between birth weight and gestational age, mediating transcripts showed limited overlap between outcomes, with most mediators being outcome-specific **(Supplementary Figure 2e)**, indicating that PFAS effects on these correlated phenotypes operate through largely distinct transcriptional mechanisms. Notably, transcripts corresponding to three genes, *TM7SF3* (transmembrane 7 superfamily member 3), *NBEAL1* (neurobeachin-like 1), and *ANXA5* (annexin A5), were significant mediators across both maternal and fetal exposure sources and both outcomes.

Functional enrichment analysis of transcriptional mediators revealed convergence on translational machinery and ribosomal components across outcomes, with shared enrichment for ’cytoplasmic translation’, ’structural constituent of ribosome’, ’cadherin binding’, and the ’Ribosome’ KEGG pathway (FDR < 0.2). Birth weight mediators showed additional enrichment for RNA processing (’ribonucleoprotein complex biogenesis’, ’RNA splicing’, ’establishment of RNA localization’), mitochondrial organization, and ubiquitin-proteasome pathways (’ubiquitin protein ligase binding’, ’Ubiquitin mediated proteolysis’), with cellular component enrichment for ’spliceosomal complex’, ’nuclear speck’, ’kinetochore’, and ’proteasome complex’. Gestational age mediators showed additional enrichment for oxidative metabolism (’aerobic respiration’, ’mitochondrial ATP synthesis coupled electron transport’) and proteostatic stress (’response to endoplasmic reticulum stress’, ’retrograde protein transport, ER to cytosol’), with molecular function enrichment for ’electron transfer activity’, ’phospholipase inhibitor activity’, and ’Hsp70 protein binding’, cellular component enrichment for ’endoplasmic reticulum protein-containing complex’, ’oxidoreductase complex’, and ’mitochondrial intermembrane space’, and KEGG enrichment for ’Protein processing in endoplasmic reticulum’ and neurodegenerative disease pathways (’Parkinson disease’, ’Huntington disease’, ’Coronavirus disease – COVID-19’).

### PFAS effects on perinatal outcomes are mediated through co-expression network hubs rather than differentially expressed transcripts

To understand whether PFAS-mediated effects on perinatal outcomes operate primarily through individual differentially expressed features or through coordinated transcriptional programs, we compared the mediation strength of differentially expressed transcripts (DETs) and genes (DEGs) against co-expression network hubs **(Figure 3a)**. This comparison tests whether PFAS effects operate primarily through individual expression changes or through coordinated perturbation of central regulatory nodes.

Across n = 144 parameter iterations **(Methods)**, we defined module hubs as the top 10 features per module per iteration ranked by absolute module eigengene correlation (|kME|), yielding a consensus set of hub features tested for mediation across all PFAS-exposure combinations. Hub transcripts that were significant mediators (n = 583) displayed markedly higher absolute ACME values compared to DETs (n = 433; Tukey HSD, p < 0.001), while hub transcripts that were non-mediators (n = 16,313) showed similar ACME distributions to DETs (p = ns) **(Figure 3d).** This finding suggests that PFAS influences birth weight primarily through perturbation of central regulatory nodes in co-expression networks rather than through transcripts showing the largest fold-changes in expression. This network-centric mediation pattern held at the gene level, where hub genes that were significant mediators (n = 75) exhibited markedly higher absolute ACME values compared to DEGs (n = 44; Tukey HSD, p < 0.001) **(Supplementary Figure 2f)**. The consistency of transcript hub-mediated effects was generalizable to gestational age as an outcome **(Supplementary Figure 2g)**, demonstrating that network-centric mediation represents a fundamental feature of PFAS effects on birth weight and gestational age.

### PFAS-mediation strength scales with transplacental transfer efficiency in outcome-specific patterns

Having determined that PFAS effects on perinatal outcomes operate primarily through transcriptional network hubs, we sought to understand whether the architecture of these effects varies systematically across compounds. PFAS compounds differ substantially in their capacity to cross the placental barrier, with transplacental transfer efficiency (TPTE), measured as the cord:maternal blood concentration ratio, ranging from < 0.5 for long-chain PFOS to > 2.0 for short-chain PFBS **(Figure 4a)**. A prerequisite for interpreting TPTE as a natural experiment is that maternal and fetal exposures reflect distinct biological signals at high TPTE rather than simply tracking one another. Consistent with this, maternal and fetal PFAS concentrations were positively correlated for low-TPTE compounds (TPTE < 1; **Supplementary Figure 3**), where limited placental transfer means fetal concentrations are largely determined by maternal levels, but uncorrelated for mid to high-TPTE compounds (TPTE > 1), where fetal concentrations become increasingly independent of maternal levels. At the transcript level, this divergence extended to differential expression: higher-TPTE compounds showed increasingly negative correlations between maternal and fetal differential expression effect estimates (r = −0.735, R^2^ = 0.540, p = 0.038), while gene-level effects showed a similar but non-significant trend (r = −0.624, R^2^ = 0.389, p = 0.098) **(Figure 4b)**, indicating that as fetal dosage increase, placental transcriptional responses to maternal versus fetal exposure become mechanistically distinct. This natural variation in the degree to which fetal and maternal exposure effects diverge provides an opportunity to test whether the underlying transcriptional mediators exhibit distinct network properties.

**Figure 4.**
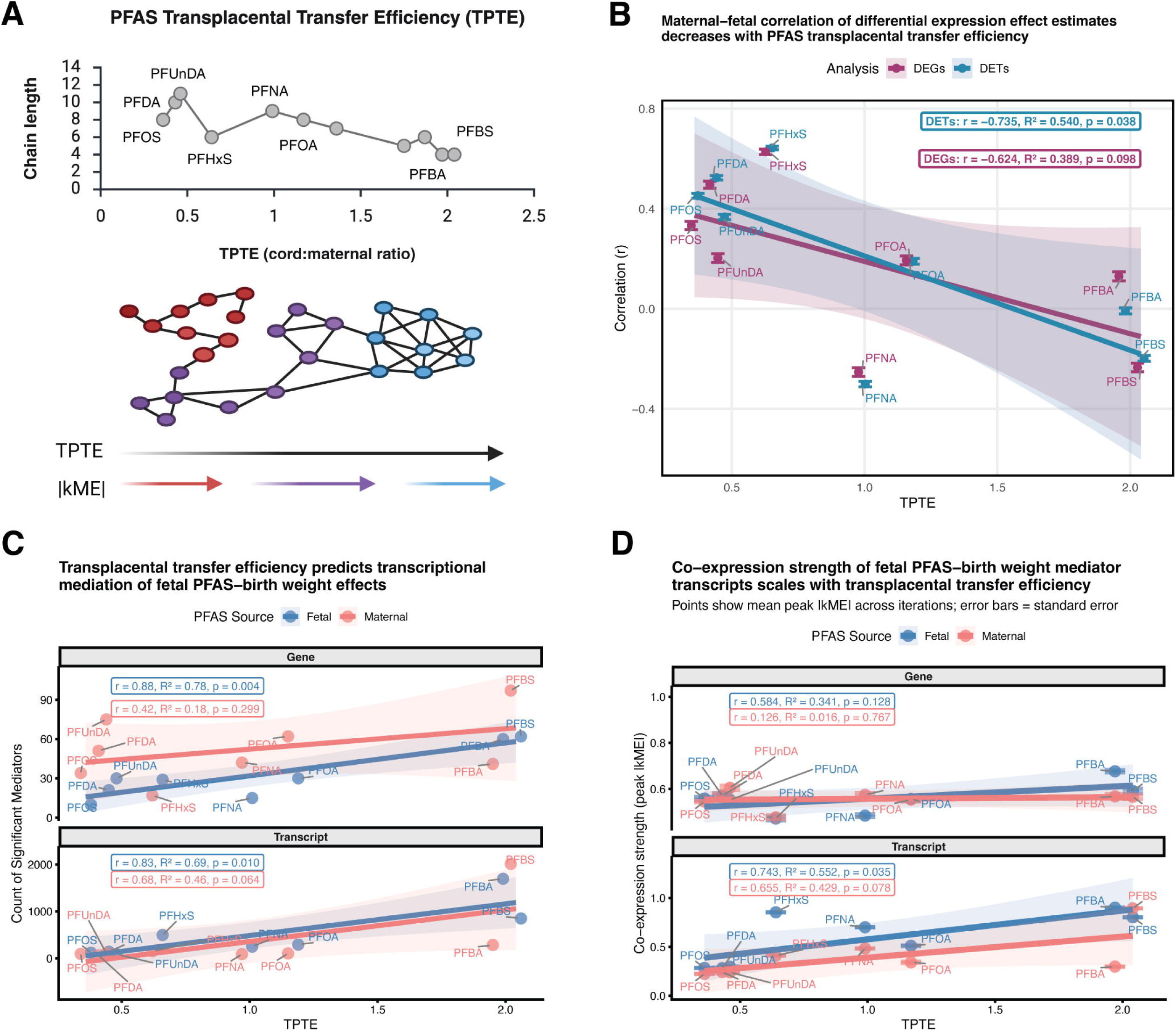
PFAS transplacental transfer efficiency (TPTE) inversely correlates with carbon chain length and associates with network-level transcriptional properties. **(A)** Top panel: TPTE, measured as the cord:maternal blood concentration ratio, increases with decreasing PFAS carbon chain length. Bottom panel: Conceptual network diagram illustrating how hub transcript module membership strength (|kME|, indicated by color intensity from red = high to blue = low) varies across PFAS compounds. Networks are shown schematically to represent the pattern whereby compounds with lower TPTE (left, red nodes) tend to have mediators with weaker module membership (lower |kME|, fewer edges), while higher TPTE compounds (right, blue nodes) have mediators with stronger co-expression relationships (higher |kME|, more edges), as quantified in subsequent panels. **(B)** Maternal-fetal correlation of transcript, but not gene-level effect estimates decrease with PFAS transplacental transfer efficiency. Pearson correlation coefficient (r) between maternal and fetal differential expression log_2_ fold changes (mean ± SE across transcripts) plotted against TPTE for each PFAS compound and analysis level. **(C)** TPTE predicts transcriptional mediation of fetal PFAS-birth weight effects. Count of significant mediators (p < 0.05, 95% CI excludes 0) across all modules and iterations plotted against TPTE for each PFAS compound and exposure source. Points represent total counts per PFAS-exposure combination. Linear regression statistics shown for fetal and maternal exposures, and for gene and transcript-level effects, separately. **(D)** Co-expression strength of fetal PFAS-birth weight mediator transcripts scales with TPTE. Peak absolute module eigengene (|kME|) values (mean ± SE across iterations) plotted against TPTE for each PFAS compound, exposure source, and analysis level. Linear regression statistics shown for fetal (blue) and maternal (pink) exposures separately.

For birth weight, both the number and co-expression strength of mediators scaled positively with TPTE at the transcript level, particularly for fetal exposures. The count of significant fetal exposure mediators increased with TPTE for both transcript-level (fetal: r = 0.83, R^2^ = 0.69, p = 0.010; maternal: r = 0.68, R^2^ = 0.46, p = 0.064) and gene-level analyses (fetal: r = 0.88, R^2^ = 0.78, p = 0.004; maternal: r = 0.42, R^2^ = 0.18, p = 0.299) **(Figure 4c)**. However, co-expression strength, measured as peak |kME| values (the mode of the kernel density distribution of absolute eigengene correlations for significant mediators, averaged across iterations), revealed transcript-specific effects: transcript-level mediators showed significant increases in |kME| with TPTE for fetal exposures (r = 0.743, R^2^ = 0.552, p = 0.035), with a similar trend for maternal exposures (r = 0.655, R^2^ = 0.429, p = 0.078), while gene-level mediators showed no significant relationship (fetal: r = 0.584, R^2^ = 0.341, p = 0.128; maternal: r = 0.126, R^2^ = 0.016, p = 0.767) **(Figure 4d)**. For gestational age, fetal mediator counts also increased with TPTE (r = 0.87, R^2^ = 0.76, p = 0.005), as did fetal co-expression strength (r = 0.904, R^2^ = 0.818, p = 0.002), mirroring the pattern observed for birth weight **(Supplementary Figure 3a–b)**. This shared fetal-specific TPTE scaling of mediator count and co-expression strength across outcomes suggests that direct fetal exposure systematically recruits more numerous and tightly co-expressed mediators regardless of the downstream perinatal outcome affected.

### TPTE reveals network compartmentalization of PFAS-mediation effects

Having established that mediator quantity and co-expression coherence scale with TPTE, we next asked whether maternal and fetal mediators occupy shared or distinct positions within co-expression networks as fetal exposure increases. We examined two complementary aspects of network topology: (1) the proximity of mediators to central regulatory hubs, quantified as edge-weighted shortest path distances, and (2) the spatial separation between fetal and maternal mediator subpopulations within shared modules **(Figure 5a)**.

**Figure 5.**
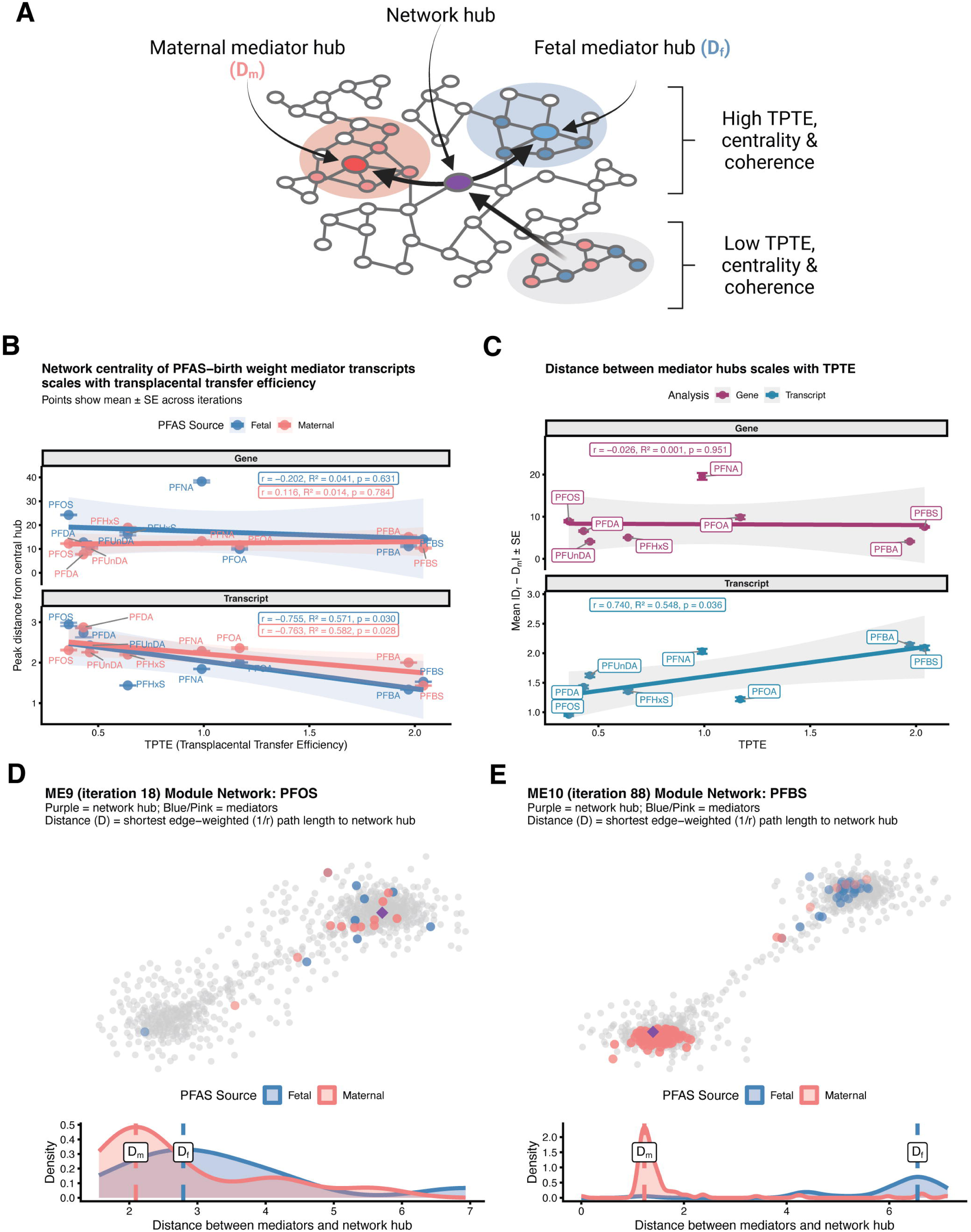
Transplacental transfer efficiency (TPTE) is associated with network compartmentalization of PFAS-mediation effects on birth weight. **(A)** Conceptual diagram illustrating the relationship between PFAS TPTE and the organization of mediator networks. For PFAS with high TPTE, mediating transcripts form distinct, well-defined hubs based on exposure source: maternal mediator hubs (D_m_, pink shaded regions) and fetal mediator hubs (D_f_, blue shaded regions). These source-specific hubs exhibit high centrality (short distances between mediators and the network hub) and coherence (functional similarity within mediator hubs). In contrast, PFAS with low TPTE show overlapping maternal and fetal mediator hubs with reduced centrality and coherence, indicating less distinct clusters (gray shaded region, lower panel). Purple node represents the network hub. **(B)** Distance between mediators and network hubs plotted against TPTE for each PFAS compound, exposure source, and analysis level. For each module-PFAS-exposure-iteration combination, peak distance is defined as the mode of the kernel density distribution of edge-weighted (1/r) shortest path lengths from the module’s central hub to all significant mediators, averaged across iterations. **(C)** Absolute difference between fetal (D_f_) and maternal (D_m_) peak distances (mean ± SE across modules and iterations) plotted against TPTE for each PFAS compound and analysis level. **(D–E)** Network diagrams (top) show module transcripts with significant mediators colored by exposure source (blue = fetal, pink = maternal; node opacity scaled by distance from hub) and the central network hub (purple diamond). Kernel density plots (bottom) show distributions of edge-weighted path lengths from hub to mediators, with dashed lines indicating peak distances (D_f_ and D_m_). **(D)** ME9 (iteration 18) Module Network with PFOS-birth weight mediator transcripts highlighted. **(E)** ME10 (iteration 88) Module Network with PFBS-birth weight mediator transcripts highlighted.

Network centrality of mediator transcripts increased with TPTE for birth weight in both exposure sources, while gene-level mediators showed systematically greater distances from hubs with no TPTE relationship. For fetal exposures at the transcript level, average distance from mediators to network hubs decreased with increasing TPTE (r = -0.755, R^2^ = 0.571, p = 0.030), with maternal exposures showing a similar relationship (r = -0.763, R^2^ = 0.582, p = 0.028) **(Figure 5b)**. In contrast, gene-level mediators exhibited distances approximately 10-fold greater than transcript-level mediators (gene-level range: ∼10-40, versus transcript-level range: ∼1-3), with no significant TPTE associations (fetal: r = -0.202, R^2^ = 0.041, p = 0.631; maternal: r = 0.116, R^2^ = 0.014, p = 0.784). This systematic difference reflects the mathematical consequence of gene-level aggregation: summing expression across isoforms dilutes pairwise correlations between co-regulated transcripts **(Supplementary Figure 4c)**, yielding weaker edge weights (lower r) and thus longer path distances (1/r) throughout the network. For gestational age, the association between TPTE and network centrality was not significant for either fetal or maternal exposures (fetal: r = −0.702, R^2^ = 0.493, p = 0.052; maternal: r = −0.389, R^2^ = 0.151, p = 0.341) **(Supplementary Figure 4d)**.

Critically, the spatial separation between fetal and maternal mediator hubs within modules exhibited TPTE-dependent organization at the transcript level but not at the gene level for birth weight. At the transcript level, the absolute difference in peak distances (|D_f_ - D_m_|) increased with TPTE (r = 0.740, R^2^ = 0.548, p = 0.036), with low-TPTE compounds like PFOS and PFDA showing minimal separation between fetal and maternal mediator hubs (|D_f_ - D_m_| ≈ 1.0-1.5), whereas high-TPTE compounds like PFBA and PFBS exhibited pronounced separation (|D_f_ - D_m_| ≈ 2.5-3.0) **(Figure 5c)**. Gene-level analyses showed greater absolute separation (|D_f_ - D_m_| range: ∼10-20), again due to aggregation-induced correlation dilution and thus a lower baseline co-expression strength, but no association with TPTE (r = -0.026, R^2^ = 0.001, p = 0.951). The lack of TPTE relationship at the gene level indicates that isoform-resolved quantification is necessary to detect the dose-dependent compartmentalization of maternal and fetal regulatory programs within co-expression networks. For gestational age, compartmentalization showed no significant association with TPTE at the transcript level (r = 0.669, R^2^ = 0.447, p = 0.070) **(Supplementary Figure 3e)**, further distinguishing the network architecture of gestational age effects from the pronounced dose-dependent compartmentalization observed for birth weight.

This TPTE-dependent spatial organization is illustrated by comparing module-level network structures for contrasting PFAS-module combinations. In module 9 (ME9; iteration 18) for PFOS exposure, a low-TPTE compound, fetal and maternal birth weight mediators showed overlapping distributions around the network hub, with minimal separation between D_f_ and D_m_ **(Figure 5d)**. In contrast, ME10 (iteration 88) for PFBS exposure, a high-TPTE compound, exhibited clear spatial separation between fetal and maternal birth weight mediators, with distinct peak distances from the central hub **(Figure 5e)**.

Together, these findings reveal a partially shared and partially outcome-specific architecture of TPTE-dependent transcriptional mediation **(Figure 6)**. Both outcomes show TPTE-dependent scaling of fetal mediator count and co-expression strength, but only birth weight exhibits the additional scaling of network centrality and compartmentalization, suggesting that fetal growth effects require coordinated, spatially organized placental-fetal regulatory programs that gestational age effects do not engage. These patterns are detectable only at the transcript level.

**Figure 6.**
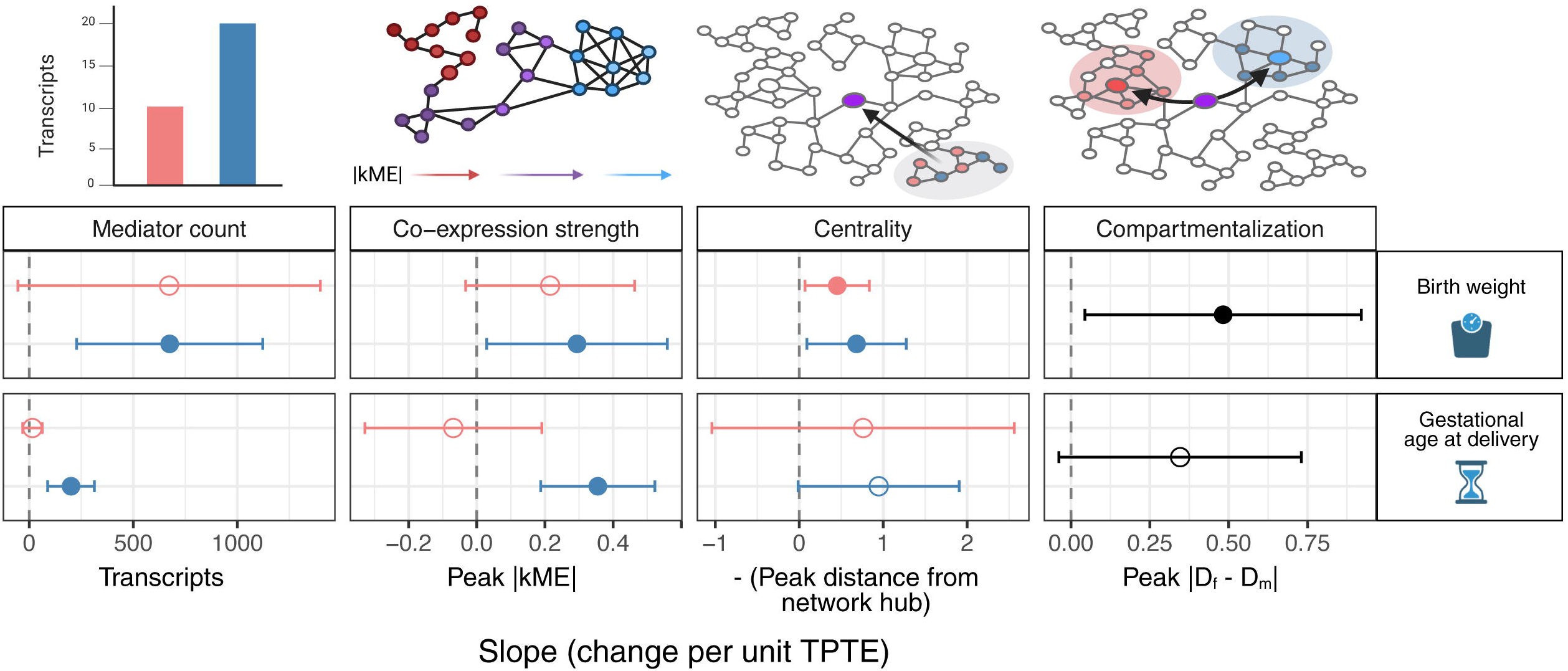
Outcome-specific scaling of transcriptional mediation properties with transplacental transfer efficiency (TPTE). Relationships between TPTE and four transcript-level mediation properties: mediator count, co-expression strength (peak |kME|), network centrality (inverse of the peak distance between mediators and network hubs), and network compartmentalization of maternal versus fetal mediators. Points represent linear regression slopes with 95% confidence intervals (error bars). Filled points indicate statistically significant associations (p < 0.05); open points indicate non-significant trends. Pink and blue distinguish exposure source (maternal versus fetal, respectively). Black is compartmentalization.

## Discussion

In this study, we leveraged isoform-resolved placental transcriptomics to elucidate the molecular mechanisms underlying PFAS effects on perinatal outcomes. A central challenge in environmental toxicology is understanding why chemically similar compounds within the same class produce heterogeneous health effects. PFAS compounds vary substantially in their capacity to cross the placental barrier,^37^ and we hypothesized that this natural variation in transplacental transfer efficiency (TPTE), reflecting different degrees of direct fetal exposure, would reveal distinct transcriptional architectures underlying compound-specific effects. Using birth weight and gestational age as complementary perinatal outcomes, we demonstrate that tissue-specific transcriptome assembly enhances detection of PFAS-responsive isoform-level changes, that PFAS effects operate primarily through co-expression network hubs rather than differentially expressed features, and that natural variation in TPTE systematically associates with mediator quantity, co-expression coherence, network centrality, and spatial organization of maternal versus fetal regulatory programs. Critically, these patterns are masked by gene-level aggregation.

### Isoform-level resolution reveals hidden transcriptional responses to PFAS exposure

The concordance between observational findings from the GUSTO prebirth study and experimental validation using patient-derived placental explants provides empirical support for tissue-specific, isoform-resolved transcriptome references in environmental exposure studies. While gene-level differential expression showed similar overlap between observational and experimental data regardless of reference annotation, isoform-level effects demonstrated substantially improved concordance using our long-read placental reference transcriptome compared to the generic GENCODE annotation. Our findings indicate that reliance on generic reference annotations fails to capture tissue-specific alternative splicing events that may be particularly susceptible to xenobiotic exposures.^21,38–40^ The substantially greater differential expression detected at the isoform level compared to gene-level aggregation aligns with emerging evidence that alternative splicing represents a critical regulatory layer in cellular responses to environmental stressors.^41,42^ Alternative splicing generates functionally distinct protein isoforms from a single gene, enabling nuanced cellular responses masked by gene-level quantification.^43^ This phenomenon may be particularly relevant for endocrine-disrupting chemicals like PFAS,^44,45^ where subtle shifts in isoform usage could alter receptor-ligand interactions or downstream signaling cascades without dramatic changes in total gene expression.^46,47^

### Network hub transcripts as primary mediators of PFAS effects on perinatal outcomes

Our finding that PFAS influences both birth weight and gestational age through perturbation of co-expression network hubs rather than differentially expressed transcripts challenges conventional assumptions in transcriptomic studies of environmental exposures. Traditional differential expression analysis prioritizes genes showing the largest fold-changes, assuming transcriptional magnitude correlates with phenotypic impact. However, our results demonstrate that transcripts with strong co-expression connectivity exert disproportionate influence on downstream phenotypes regardless of their individual expression changes. Hub transcripts occupy critical regulatory positions within transcriptional programs, enabling them to coordinate responses across multiple biological pathways.^28,48^ Perturbation of these regulatory nodes may propagate effects throughout co-expression networks, amplifying relatively modest changes in individual transcript levels into substantial phenotypic consequences.^49^

This divergence between hub-mediated effects and differential expression reflects fundamental differences in how gene-level versus transcript-level features participate in regulatory networks. Co-expression networks are defined by pairwise correlations between features, and network distances are calculated as the inverse of correlation strength (1/r). When multiple isoforms from the same gene respond differently to PFAS exposure—for example, because specific isoforms are preferentially expressed in distinct placental cell types, or because alternative splicing is itself PFAS-responsive—summing expression across all isoforms dilutes these specific regulatory relationships. Weaker correlations throughout gene-level networks yield systematically longer path distances, reducing the apparent connectivity of regulatory features and obscuring the coordinated responses that define hub-mediated effects. In contrast, transcript-level networks capture isoform-specific co-regulation, revealing that hub transcripts, despite often showing modest individual expression changes, mediate PFAS effects more strongly than differentially expressed features precisely because of their coordinated regulatory relationships.

The generalizability of network-centric mediation across both birth weight and gestational age suggests that hub-based mediation represents a general principle of how environmental exposures affect complex perinatal phenotypes.^32,33,50^ Functional enrichment analysis revealed that PFAS compounds perturb fundamental cellular homeostatic processes including RNA processing, protein degradation, and translational machinery, with outcome-specific effects manifesting through differential engagement of pathways. Birth weight mediators emphasized intracellular trafficking and quality control, consistent with placental nutrient transport and cellular growth regulation, while gestational age mediators were enriched for oxidative phosphorylation and mitochondrial function, reflecting the high energetic demands of parturition signaling. The recurrent enrichment of neurodegenerative disease pathways likely reflects shared cellular stress responses including protein aggregation, mitochondrial dysfunction, and oxidative stress, rather than direct neurological effects in placental tissue.^51,52^

While most mediators were exposure source and outcome-specific, transcripts corresponding to three genes (*TM7SF3*, *NBEAL1*, and *ANXA5*) represent the only transcripts showing consistent mediation across all four exposure–outcome combinations tested. These three genes implicate distinct but complementary mechanisms that collectively span the major pathways through which PFAS are understood to perturb placental biology. *TM7SF3* encodes a seven-transmembrane protein that attenuates endoplasmic reticulum (ER) stress and suppresses the unfolded protein response (UPR),^53^ while also localizing to nuclear speckles to regulate alternative splicing of hundreds of genes.^54^ *TM7SF3* expression dysregulation tips trophoblasts toward apoptosis and impairs the fine balance of ER stress required for normal syncytialization and placental growth.^55,56^ *NBEAL1* is a Golgi-associated regulator of *SREBP2* processing that controls LDL receptor expression and downstream cholesterol homeostasis.^57^ PFAS-driven dysregulation of the *SREBP2* axis in trophoblasts may impair the cholesterol supply to the fetus required for growth and steroid hormone synthesis.^58,59^ Finally, *ANXA5*, originally discovered from human placenta,^60^ encodes annexin A5, a calcium-dependent phospholipid-binding protein highly expressed on the syncytiotrophoblast surface where it provides anticoagulant protection by shielding phosphatidylserine-exposing membranes,^61^ and whose 2D-lattice assembly mediates cytotrophoblast fusion.^62^ Reduced *ANXA5* expression is a consistent finding in placentas from pregnancies complicated by intrauterine growth restriction and preeclampsia.^62,63^ The functional enrichment of gestational age mediators for ER stress response and phospholipase inhibitor activity, together with the identification of *TM7SF3* and *ANXA5* as consistent mediators, suggests that PFAS-induced proteostatic and membrane stress are associated with both fetal growth restriction and altered parturition timing through placental transcriptional mechanisms.

### Natural variation in TPTE reveals isoform-specific transcriptional architectures

TPTE reflects the net permeability of the placenta to individual PFAS compounds, governed by membrane transporter-mediated uptake into the syncytiotrophoblast and compound-specific partitioning that determines efflux to the fetal circulation. The systematic association between TPTE and multiple dimensions of placental transcriptional network architecture, detectable only at the transcript level, represents our most striking finding, with the pattern of scaling differing instructively between outcomes **(Figure 6)**. For both birth weight and gestational age, direct fetal tissue exposure engages dose-dependent recruitment of increasingly numerous and synchronized transcriptional mediators, suggesting that as more compound reaches fetal tissues, more extensive and coherent placental transcriptional responses are mobilized regardless of the downstream perinatal outcome. However, the two outcomes diverge in network architecture. For birth weight, the steep positive scaling of all network properties with TPTE suggests that fetal growth effects operate through coordinated placental-fetal metabolic programs, including insulin-like growth factor signaling, amino acid transport, and mitochondrial biogenesis,^64–66^ where higher-TPTE compounds elicit increasingly synchronized and spatially segregated transcriptional responses. In contrast, the absence of TPTE-dependent network centralization and compartmentalization for gestational age is consistent with parturition timing being governed by predominantly maternal physiological cascades^67^ in which the degree of fetal exposure has limited influence on network architectural organization, even as mediator count and co-expression strength respond to fetal dose.

### Limitations and future directions

Several limitations merit consideration. Our observational design cannot definitively establish causal relationships, though experimental validation using patient-derived explants provides supporting evidence. Our findings highlight two complementary requirements for advancing causal inference in environmental genomics. First, the underlying molecular measurements must be well-calibrated. We demonstrate that isoform-resolved transcriptome references improve concordance between observational and experimental effect estimates, providing a foundation for more reliable causal inference. Second, the mediation analysis methods themselves require further development to address unmeasured confounding. The standard mediation framework assumes no unmeasured confounders conditional on measured covariates, an assumption that cannot be empirically verified in observational prebirth studies.^68–72^ While we leverage TPTE as a natural experiment to partially address this limitation, future work employing negative control approaches, such as using maternal epigenomic data as proxies for unmeasured confounders^73–75^, will be essential for strengthening causal claims. This two-pronged approach improves both the molecular measurements and the causal inference methods and represents a path forward for developmental toxicology studies.

The biological interpretation of outcome-specific TPTE-dependent patterns remains speculative; the divergent network architectures may reflect differences between fetal-mediated versus maternal-mediated mechanisms, but alternative explanations, including differences in measurement error, statistical power, or unmeasured confounding, cannot be excluded. Moreover, our analysis focused on term placentas with East/Southeast/South Asian ancestry; validation in diverse populations is necessary to establish generalizability. Additionally, bulk tissue analysis obscures cell-type-specific responses. Single-cell or spatial transcriptomics will be essential for determining whether PFAS effects concentrate in specific cell populations.^76^ Furthermore, we examined only eight PFAS compounds, whereas hundreds of PFAS are in commercial use and detected in human biomonitoring studies, including emerging short-chain replacements whose toxicological profiles remain poorly characterized.^2^ Whether the TPTE-dependent transcriptional architectures we identify generalize across the broader PFAS landscape, particularly to compounds with distinct functional groups or branched-chain structures, remains to be determined. Finally, while our network-based approach revealed that hub transcripts mediate PFAS effects, experimental perturbation of candidate hub transcripts in placental cell cultures or organoid models will be essential for testing causal mechanisms linking network position to phenotypic impact.

### Implications for developmental toxicity studies

Three broad principles emerge from this work: gene-level aggregation obscures regulatory relationships through which environmental exposures perturb developmental processes; environmental toxicants operate through central regulatory nodes rather than dramatic transcriptional changes; and natural variation in transplacental transfer efficiency systematically associates with transcriptional network architecture, providing a generalizable framework for resolving mechanistic heterogeneity that may extend beyond PFAS to other placental-crossing contaminants. This framework integrates advances in molecular measurement with evolving causal inference methodologies. The improved concordance between observational and experimental effect estimates with tissue-specific isoform resolution demonstrates that measurement quality directly translates to more reliable mechanistic inference, and combined with emerging negative control methods for unmeasured confounding, offers a path toward resolving compound-specific mechanisms from bulk tissue assays across environmental contaminant classes. For risk assessment, compound-specific transfer efficiency should be considered alongside traditional toxicity metrics, as compounds achieving similar maternal exposures may exert vastly different effects depending on fetal tissue concentrations.^77^ Regulatory frameworks based solely on maternal exposure levels may systematically underestimate risks from high-TPTE compounds.^78^ Most broadly, natural variation in toxicokinetic properties presents an underutilized resource for dissecting the molecular mechanisms underlying heterogeneous environmental health effects.

## Materials and Methods

### GUSTO prebirth study biological samples, PFAS measurements, and placental RNA sequencing

We included placental villous tissue samples from term, live births without known placentation pathologies from the Growing Up in Singapore Toward Healthy Outcomes (GUSTO) prebirth study.^17,35^ Of n = 783 umbilical cord plasma samples analyzed for PFAS in GUSTO, we utilized n = 124 samples with available placental RNA-seq data and PFAS measurements in maternal blood (n = 101) and cord blood (n = 101) at delivery (n = 78 with paired maternal and cord blood PFAS measurements).^77,79^ For gestational age analyses, samples were restricted to births following spontaneous onset of labor (regardless of mode of delivery; n = 72 with any PFAS measure; n = 42 with paired maternal and cord blood PFAS measurements). Gestational age was based on a first trimester ultrasound scan of fetal crown rump length.

Maternal samples were collected when women were admitted to the National University Hospital Singapore for delivery. Umbilical cord venous blood was collected immediately following delivery into EDTA tubes. Samples were centrifuged at 1,600 g for 10 min at 4 °C to isolate plasma, then further centrifuged at 16,000 g for 10 min at 4 °C. Four mL of plasma was added to a plain tube containing ∼15 µL trasylol and stored at −80 °C in 0.4 mL aliquots until batch analysis. PFAS were analyzed using ultra-performance liquid chromatography-tandem mass spectrometry (UPLC-MS/MS). Eight PFAS were measured: perfluorobutanoic acid (PFBA), perfluoroundecanoic acid (PFUnDA), perfluorobutanesulfonic acid (PFBS), perfluorooctanoic acid (PFOA), perfluorononanoic acid (PFNA), perfluorodecanoic acid (PFDA), perfluorohexanesulphonic acid (PFHxS), and perfluorooctanesulfonic acid (PFOS). All measured PFAS had detection rates ≥82%. Values below the limit of quantification (LOQ; signal-to-noise ratio < 10) and limit of detection (LOD; signal-to-noise ratio < 3) were imputed with LOQ/2 and LOD/2, respectively.^77,79^

Placenta samples were processed within 1 hour of delivery. Five villous biopsies of each placenta were rinsed with phosphate buffer saline (PBS), snap-frozen with liquid nitrogen and stored at −80 °C before RNA extraction. RNA was extracted from approximately 100 mg of crushed tissue using phenol-chloroform and purified with a Qiagen RNeasy Mini Kit. Paired-end Illumina libraries were sequenced as described previously.^80,81^ RNA quality was assessed by Nanodrop spectrophotometry and Agilent TapeStation, with ribosomal RNA depletion performed using Illumina Ribo-Zero kits. Sequencing libraries were prepared using NEB Next Ultra Directional RNA Library Prep Kit and sequenced on Illumina HiSeq 4000 platform to a minimum of 50 M paired-end 150 bp reads per library.

### In vitro experimental explant tissue procurement and culture

Placental villous tissues (n = 3 term and n = 1 second trimester) were obtained fresh from non-laboring patients at UC San Diego (UCSD) Health-affiliated hospital/clinics and transported on ice in RPMI media to the Elkin lab for processing. Chorionic villi were dissected into five approximately 5 mg pieces per sample, yielding five explants totaling 20-30 mg per sample. Explants were cultured in RPMI 1640 medium supplemented with 10% FBS and 1% P/S at 37 °C in a 5% CO_2_ humidified incubator. Samples were placed in 12-well tissue culture plates and exposed to three PFBS concentrations: 0 µM (control), 5 µM, and 20 µM. Each treatment was performed in technical triplicate, with one plate per sample. Prior to each experiment, a 1 M PFBS stock solution was thawed to room temperature and diluted to 1 mM in supplemented RPMI 1640 medium. Working concentrations of 5 µM and 20 µM were then prepared from this intermediate stock.

After the 24-hour treatment period, explants from each sample were transferred to 2 mL tubes prefilled with 2.8 mm stainless steel beads in 700 µL of QIAzol® Lysis Reagent. Samples were homogenized using a Benchmark BeadBug Homogenizer at 3000 rpm for 6 cycles of 30 seconds each, followed by 3 additional cycles at 2500 rpm. mRNA was extracted using the QIAGEN miRNeasy® Mini Kit according to the manufacturer’s protocol and eluted in RNase-free water. RNA concentration and purity were quantified by NanoDrop (Thermo Fisher Scientific), and samples were stored at -20 °C until further processing. For term samples, each treatment group was performed in biological triplicate (three separate villous samples from the same placenta) within each experiment. RNA from these triplicates was pooled by treatment group prior to sequencing, yielding nine term libraries representing three placentas × three treatment groups (0 µM control, 5 µM, and 20 µM PFBS). For the second-trimester sample, nine libraries were sequenced from a single placenta: three treatment groups each performed in technical triplicate, yielding nine libraries. In total, 18 libraries were generated from 4 placentas (3 term, 1 second trimester). Pooled RNA samples were sent to Novogene for sequencing. Poly(A)-enriched libraries were prepared and sequenced as unstranded cDNA on the NovaSeq X Plus platform using paired-end 150 bp Illumina chemistry.

### Placental transcriptome assembly and annotation

We previously assembled an isoform-resolved placental transcriptome reference^36^ from long-read Oxford Nanopore sequencing of n = 72 term placental samples from live births without known placentation pathologies. Samples included n = 6 from the GUSTO prebirth study cohort and n = 66 from an independent National University Hospital Singapore prebirth study cohort.^81^ Placentae were processed within 1 hour of delivery, with five villous biopsies per placenta rinsed with PBS, snap-frozen, and stored at -80 °C. RNA was extracted using phenol-chloroform and samples were sequenced on a PromethION platform with PCR-cDNA library preparation. FAST5 files were base-called with Guppy (GPB_2003_v1_revAX_14Dec2018), and full-length cDNA reads were identified with Pychopper2,^82^ error-corrected with FMLRC2,^83^ and aligned to hg38 using Minimap2.^84^ De novo transcript assembly was performed with ESPRESSO.^85^ Assembled transcripts were classified relative to GENCODE v45 using SQANTI3.^86^ High-confidence isoforms required minimum coverage of 3 long reads per transcript and 3 tissue-matched Illumina short reads per splice junction, with transcription start and termination sites validated against CAGE-seq, DNase-seq, and polyA site datasets. Additional filters removed assembly artifacts including reverse transcriptase switching and intrapriming.

### Quantification of gene and isoform expression

We used fastp^87^ to trim sequencing adapters and filter low-quality reads. We quantified short-read RNA-seq data from GUSTO samples against both our isoform-resolved placental transcriptome reference (lr-assembly) and the GENCODEv45 transcriptome reference. Decoy-aware transcriptome indices for Salmon^88^ were constructed with MashMap.^89^ Transcriptome quantification was then performed with Salmon with settings --libType A --validateMappings --seqBias and including 50 bootstrap replicates. Transcript quantifications were loaded into R with tximport,^43^ and overdispersion arising from read-to-transcript mapping ambiguity was estimated with catchSalmon() from the edgeR package^90^ and divided from the counts prior to downstream analyses.^91^ To reduce noise in downstream analyses, we filtered the expression matrices to retain only transcripts with at least 5 read counts in a minimum of 3 samples.

### Weighted co-expression network analyses

To identify co-expressed transcript modules associated with PFAS exposure, we performed weighted co-expression network analysis using the WGCNA package.^92^ Transcript or gene expression data were first filtered to retain features detected in at least 20% of samples with variance above the 10th percentile. Count data were normalized using variance stabilizing transformation (VST) via DESeq2.^93^ We applied surrogate variable analysis (SVA) with RUVr^94^ to identify and regress out one latent factor of technical variation while preserving variation associated with PFAS exposures. Batch effects from the surrogate variable were removed using removeBatchEffect from the limma package.^95^ Signed co-expression networks were constructed using the WGCNA package methodology applied to transcript- and gene-level expression across a systematic parameter space to ensure robustness. We performed 144 iterations spanning combinations of soft-thresholding power (transcript-level analysis: 3, 4, 5; gene-level analysis: 14, 15, 16), minimum module size (30, 50, 75, 100), tree-cutting depth (deepSplit: 2, 3, 4), and module merging threshold (mergeCutHeight: 0.15, 0.20, 0.25, 0.30). Soft-thresholding powers were selected as the first three integer values achieving scale-free topology fit (R^2^) ≥ 0.85, ensuring networks approximate scale-free architecture while minimizing information loss from excessive thresholding. For each iteration, modules were identified via hierarchical clustering of topological overlap matrix (TOM) dissimilarity, followed by dynamic tree cutting and eigengene-based merging of similar modules. Module hub transcripts were defined as the top 10 features per module per iteration ranked by absolute module eigengene correlation (|kME|), yielding a consensus set of hub features across the parameter space. For each module, the module eigengene was calculated as the first principal component of module expression, and kME values were computed as Pearson correlations between individual feature expression and the module eigengene.

### Differential expression and mediation analyses

Differential transcript and gene expression analyses were performed with DESeq2. Models adjusted for gestational age (weeks) z-score, fetal sex, and *V* dimensions of technical variation per PFAS compound × exposure source estimated with RUVr. Features were considered significant given Benjamini-Hochberg adjusted p-value < 0.1 and absolute log_2_ fold change > 1. To assess concordance between explant and tissue PFBS effects, features were considered concordant if nominally significant (p < 0.05) in the same direction in both datasets. Enrichment of concordant features among those nominally significant in explants (|log_2_FC| > 1, p < 0.05) was tested using Fisher’s exact test, with all detected features as the universe. For DTE, concordance and enrichment were assessed at the gene level, counting genes with at least one differentially used transcript meeting criteria in both datasets.

Mediation analyses were performed for all expressed features for each compound × exposure source combination using the mediation package. We used variance stabilizing transformation (VST) normalized counts to estimate the indirect effect of PFAS exposure z-score on birth weight (grams) z-score and gestational age (weeks) z-score, respectively. Mediators were considered significant given Benjamini-Hochberg adjusted p-value < 0.05 and 95% confidence intervals excluded zero. To compare mediation effect sizes between differentially expressed transcripts (DETs) or genes (DEGs) and module hub transcripts, we compiled absolute ACME values for three categories: DETs or DEGs (FDR < 0.1, |log_2_FC| > 1), hub mediators (top 10 features per module per iteration by |kME| with significant mediation), and hub non-mediators (top 10 features per module per iteration by |kME| without significant mediation). Differences in |ACME| distributions across categories were assessed using ANOVA followed by Tukey’s HSD post-hoc test, with significance determined by bootstrapping (1000 iterations, n = 100 per category per iteration) and averaging p-values across bootstrap samples.

### Network topology and distance analyses

To assess network proximity of mediators to module hubs, we constructed weighted adjacency matrices using signed co-expression networks with soft-thresholding powers selected as the first three integer values achieving scale-free topology fit (R^2^) ≥ 0.85 (transcript-level: 3, 4, 5; gene-level: 14, 15, 16), following standard methods for WGCNA network construction. Adjacency matrices were constructed for all samples (birth weight analyses) and those from births following spontaneous onset of labor (regardless of mode of delivery; gestational age analyses) separately. Adjacency values below 0.1 were set to zero to remove weak connections. For each module within each WGCNA iteration, the network hub was identified as the feature with maximum eigenvector centrality based on edge weights. Edge-weighted shortest path distances from the hub to all significant mediators were calculated using Dijkstra’s algorithm implemented in the igraph package,^96^ with the inverse of correlation strengths (1/*r*) as edge weights. For each PFAS-exposure-module-iteration combination, the peak distance was defined as the mode of the kernel density distribution of path lengths (D_f_ for fetal exposure mediators and D_m_ for maternal exposure mediators). To obtain robust estimates, distances were averaged across WGCNA iterations for each PFAS-exposure combination. Spatial separation between fetal and maternal mediator hubs was quantified as the absolute difference between the mode of the kernel density distribution of path lengths for fetal versus maternal mediators, i.e., |D_f_ - D_m_|, for each PFAS compound, computed per iteration and then averaged across modules and iterations.

### Transplacental transfer efficiency analyses

Transplacental transfer efficiency (TPTE) was defined as the average ratio of cord blood to maternal blood PFAS concentrations within the full GUSTO prebirth study (previously estimated by Leader *et al*).^79^ Relationships between TPTE and transcriptional outcomes were assessed using Pearson correlation and linear regression on PFAS-level summary statistics (mean ± standard error across WGCNA iterations). For co-expression strength analysis, peak |kME| values were defined as the mode of the kernel density distribution of absolute eigengene correlations for significant mediators within each PFAS compound × exposure source × iteration combination, then averaged across iterations to yield mean ± SE per PFAS × exposure combination. For network centrality analysis, peak distance values were defined as the mode of the kernel density distribution of edge-weighted (1/r) shortest path lengths from the module’s central hub transcript to all significant mediators for each module × PFAS × exposure × iteration combination, then averaged across modules and iterations to yield mean ± SE per PFAS × exposure combination. Correlation of maternal and fetal differential expression effect estimates was calculated as the Pearson correlation coefficient of log_2_ fold changes between matched features for each PFAS compound.

### Functional enrichment analyses

Gene Ontology (GO) and Kyoto Encyclopedia of Genes and Genomes (KEGG) enrichment analyses were performed using the clusterProfiler package.^97^ Transcript identifiers for significant mediators were mapped to gene identifiers using a transcript-to-gene mapping table, with Ensembl gene IDs converted to Entrez Gene IDs using the org.Hs.eg.db annotation package.^98^ GO enrichment analyses were conducted separately for three ontologies: Biological Process (BP), Molecular Function (MF), and Cellular Component (CC). KEGG pathway enrichment was performed using human pathway annotations (organism code: ’hsa’). All enrichment tests applied hypergeometric testing with Benjamini-Hochberg multiple testing correction, with terms considered significantly enriched at p-value < 0.05 and FDR < 0.2. To reduce redundancy in GO results, semantic similarity-based simplification was applied using the simplify function from clusterProfiler, which removes redundant terms based on Wang’s method for measuring GO semantic similarity. KEGG results were converted to gene symbols for interpretability using the setReadable function.

## Ethics Approval

All study protocols follow the principles of the Declaration of Helsinki. The GUSTO prebirth study was approved by the National Healthcare Group Domain Specific Review Board and the SingHealth Centralized Institutional Review Board. Informed written consent was obtained from all mothers at recruitment. Procurement of the *in vitro* placental tissues was approved by University of San Diego (UCSD) Institutional Review Board (Project #181917 for term placental tissues and Project #172111 for second trimester placental tissues). Written and informed consent of the mothers was obtained in collaboration with the UCSD Center for Perinatal Discovery. All data presented in this manuscript and associated materials are published in accordance with the terms of these approvals and the informed consent obtained from study participants.

## Acknowledgements

The authors would like to thank the GUSTO prebirth study participants and their families, and the related staff and study teams, without whom this study would not be possible. We also thank the UCSD Center for Perinatal Discovery for support with tissue procurement. Additionally, we thank members of both the Bhattacharya, Huang, and Elkin labs for critical reading of our manuscript, and members of Michael I. Love’s lab for feedback on methods. GUSTO is supported by the National Research Foundation (NRF) under its Translational and Clinical Research (TCR) Flagship Programme (NMRC/TCR/004-NUS/2008 and NMRC/TCR/012-NUHS/2014 to S.-Y.C.) and the Open Fund-Large Collaborative Grant (MOH-000504 to S.-Y.C.) administered by the Singapore Ministry of Health’s National Medical Research Council (NMRC) and the Agency for Science, Technology and Research (A*STAR). Additional funding is provided by the Institute for Human Development and Potential (IHDP), A*STAR. Placental long-read RNA sequencing was supported by an NMRC Open Fund-Young Individual Research Grant (MOH-000550-00 to J.Y.H.). J.Y.H. was also supported by an A*STAR Human Health and Potential–Prenatal/Early Childhood Grant (H24P2M0002) and Pilot Projects Program funding from the NIMHD RCMI-CC (U24MD015970). E.R.E. was supported by Pilot Projects Program funding from the NIMHD RCMI-CC (U54MD012397) and the National Institute of Environmental Health Sciences, National Institutes of Health (K01ES036552 to E.R.E).

## Author Contributions

S.T.B., J.Y.H, A.B., and E.R.E. designed the study. S.-Y.C., H.E.J.Y., and J.K.Y.C. managed the GUSTO study. E.R.E, M.G.D., S.N.C., A.E.T., G.R.R., and C.M.C. performed the *in vitro* experiments with patient-derived placental explants. S.T.B. directed bioinformatic and statistical analyses and wrote the initial draft of the manuscript. S.L. contributed to bioinformatic and statistical analyses. All authors contributed to the writing and editing of the final manuscript.

## Data Availability

Datasets required to reproduce the analyses conducted in this study are available in the associated Supplementary Data or have been deposited in a GitHub repository under https://github.com/sbresnahan/placenta_pfas_networks. Illumina short reads generated through the GUSTO prebirth study, along with the associated sample covariates, are available under restricted access. Data access is governed by applicable local laws and policies. Please see https://gustodatavault.sg/about/request-for-data for details on procedures for requesting access to GUSTO data.

## References

1. Fenton, S. E. et al. Per-and polyfluoroalkyl substance toxicity and human health review: current state of knowledge and strategies for informing future research. Environmental Toxicology and Chemistry 40, 606–630 (2021).

2. Glüge, J. et al. An overview of the uses of per-and polyfluoroalkyl substances (PFAS). Environmental Science: Processes & Impacts 22, 2345–2373 (2020).

3. Sunderland, E. M. et al. A review of the pathways of human exposure to poly-and perfluoroalkyl substances (PFASs) and present understanding of health effects. Journal of Exposure Science & Environmental Epidemiology 29, 131–147 (2019).

4. Botelho, J. C., Kato, K., Wong, L.-Y. & Calafat, A. M. Per- and polyfluoroalkyl substances (PFAS) exposure in the U.S. population: NHANES 1999–March 2020. Environmental Research 270, 120916 (2025).

5. DeLuca, N. M. et al. Geographic and demographic variability in serum PFAS concentrations for pregnant women in the United States. J Expo Sci Environ Epidemiol 33, 710–724 (2023).

6. Bline, A. P. et al. Public Health Risks of PFAS-Related Immunotoxicity Are Real. Curr Environ Health Rep 11, 118–127 (2024).

7. Mamsen, L. S. et al. Concentrations of perfluoroalkyl substances (PFASs) in human embryonic and fetal organs from first, second, and third trimester pregnancies. Environment International 124, 482–492 (2019).

8. Wu, M. et al. Prenatal per- and polyfluoroalkyl substances (PFAS) exposure and maternal thyroid homeostasis: Nonlinear, compound-specific, and mixture effects. Environmental Chemistry and Ecotoxicology 7, 1280–1288 (2025).

9. McAdam, J. & Bell, E. M. Determinants of maternal and neonatal PFAS concentrations: a review. Environ Health 22, 41 (2023).

10. Abdulkadir, A. et al. Epigenetic Consequences of In Utero PFAS Exposure: Implications for Development and Long-Term Health. Int J Environ Res Public Health 22, 917 (2025).

11. Verner, M.-A. et al. Associations of Perfluoroalkyl Substances (PFAS) with Lower Birth Weight: An Evaluation of Potential Confounding by Glomerular Filtration Rate Using a Physiologically Based Pharmacokinetic Model (PBPK). Environ Health Perspect 123, 1317–1324 (2015).

12. Wright, J. M. et al. Birth weight in relation to maternal and neonatal biomarker concentration of perfluorooctane sulfonic acid: a meta-analysis and meta-regression from a systematic review. J Expo Sci Environ Epidemiol 35, 1030–1040 (2025).

13. Starling, A. P. et al. Prenatal exposure to per-and polyfluoroalkyl substances and maternal and neonatal thyroid function in the Healthy Start Study: A prospective cohort study. Thyroid 31, 1066–1078 (2021).

14. Szilagyi, J. T., Avula, V. & Fry, R. C. Perfluoroalkyl substances (PFAS) and their effects on the placenta, pregnancy and child development: A potential mechanistic role for placental peroxisome proliferator-activated receptors (PPARs). Curr Environ Health Rep 7, 222–230 (2020).

15. Furness, A. I. Maternal-fetal conflict and the timing of birth. *Evolution*, Medicine, and Public Health 13, 292–306 (2025).

16. Burton, G. J. & Fowden, A. L. The placenta: a multifaceted, transient organ. Philos Trans R Soc Lond B Biol Sci 370, 20140066 (2015).

17. Chen, L.-W. et al. Associations of cord plasma per- and polyfluoroakyl substances (PFAS) with neonatal and child body composition and adiposity: The GUSTO study. Environment International 183, 108340 (2024).

18. Norén, E. et al. Transplacental transfer efficiency of perfluoroalkyl substances (PFAS) after long-term exposure to highly contaminated drinking water: a study in the Ronneby Mother-Child Cohort. J Expo Sci Environ Epidemiol 35, 445–453 (2025).

19. Beesoon, S. et al. Isomer-specific binding affinity of perfluorooctanesulfonate (PFOS) and perfluorooctanoate (PFOA) to serum proteins. Environmental Science & Technology 45, 7999–8004 (2011).

20. Bangma, J. T. et al. A review of recent literature on the relationship between prenatal exposure to per-and polyfluoroalkyl substances and birth outcomes. Current Environmental Health Reports 7, 338–349 (2020).

21. Annalora, A. J., Coburn, J. L., Jozic, A., Iversen, P. L. & Marcus, C. B. Global transcriptome modulation by xenobiotics: the role of alternative splicing in adaptive responses to chemical exposures. Hum Genomics 18, 127 (2024).

22. Bhattacharya, A. et al. Isoform-level transcriptome-wide association uncovers genetic risk mechanisms for neuropsychiatric disorders in the human brain. Nat Genet 55, 2117–2128 (2023).

23. Aguzzoli Heberle, B., et al. Mapping medically relevant RNA isoform diversity in the aged human frontal cortex with deep long-read RNA-seq. Nat Biotechnol 43, 635–646 (2025).

24. Wen, C. et al. Cross-ancestry atlas of gene, isoform, and splicing regulation in the developing human brain. Science 384, eadh0829 (2024).

25. Mostafavi, H., Spence, J. P., Naqvi, S. & Pritchard, J. K. Systematic differences in discovery of genetic effects on gene expression and complex traits. Nat Genet 55, 1866–1875 (2023).

26. Rice, A. M. & McLysaght, A. Dosage sensitivity is a major determinant of human copy number variant pathogenicity. Nat Commun 8, 14366 (2017).

27. Jeong, H., Mason, S. P., Barabási, A.-L. & Oltvai, Z. N. Lethality and centrality in protein networks. Nature 411, 41–42 (2001).

28. Barabási, A.-L. & Oltvai, Z. N. Network biology: understanding the cell’s functional organization. Nat Rev Genet 5, 101–113 (2004).

29. Seo, C. H., Kim, J.-R., Kim, M.-S. & Cho, K.-H. Hub genes with positive feedbacks function as master switches in developmental gene regulatory networks. Bioinformatics 25, 1898–1904 (2009).

30. Maertens, A., Tran, V., Kleensang, A. & Hartung, T. Weighted Gene Correlation Network Analysis (WGCNA) Reveals Novel Transcription Factors Associated With Bisphenol A Dose-Response. Front. Genet. 9, 508 (2018).

31. Sturla, S. J. et al. Systems Toxicology: From Basic Research to Risk Assessment. Chem. Res. Toxicol. 27, 314–329 (2014).

32. Cox, B. et al. A Co-expression Analysis of the Placental Transcriptome in Association With Maternal Pre-pregnancy BMI and Newborn Birth Weight. Front. Genet. 10, 354 (2019).

33. Deyssenroth, M. A. et al. Whole-transcriptome analysis delineates the human placenta gene network and its associations with fetal growth. BMC Genomics 18, 520 (2017).

34. Parenti, M. et al. The placental transcriptome serves as a mechanistic link between prenatal phthalate exposure and placental efficiency. Environment International 206, 109949 (2025).

35. Soh, S.-E. et al. Cohort profile: Growing Up in Singapore Towards healthy Outcomes (GUSTO) birth cohort study. Int J Epidemiol 43, 1401–1409 (2014).

36. Bresnahan, S. T. et al. Long-read transcriptome assembly reveals vast isoform diversity in the placenta associated with metabolic and endocrine function. Preprint at 10.1101/2025.06.26.661362 (2025).

37. Appel, M. et al. The transplacental transfer efficiency of per- and polyfluoroalkyl substances (PFAS): a first meta-analysis. *Journal of Toxicology and Environmental Health*, Part B 25, 23–42 (2022).

38. Lapehn, S. et al. A transcriptomic comparison of in vitro models of the human placenta. Placenta 159, 52–61 (2025).

39. Zaharieva, E., Chipman, J. K. & Soller, M. Alternative splicing interference by xenobiotics. Toxicology 296, 1–12 (2012).

40. Annalora, A. J., Marcus, C. B. & Iversen, P. L. Alternative Splicing in the Nuclear Receptor Superfamily Expands Gene Function to Refine Endo-Xenobiotic Metabolism. Drug Metabolism and Disposition 48, 272–287 (2020).

41. Biamonti, G. & Caceres, J. F. Cellular stress and RNA splicing. Trends in Biochemical Sciences 34, 146–153 (2009).

42. Laloum, T., Martín, G. & Duque, P. Alternative Splicing Control of Abiotic Stress Responses. Trends in Plant Science 23, 140–150 (2018).

43. Soneson, C., Love, M. I. & Robinson, M. D. Differential analyses for RNA-seq: transcript-level estimates improve gene-level inferences. F1000Res 4, 1521 (2016).

44. Coperchini, F. et al. Thyroid Disrupting Effects of Old and New Generation PFAS. Front. Endocrinol. 11, 612320 (2021).

45. Mokra, K. Endocrine Disruptor Potential of Short- and Long-Chain Perfluoroalkyl Substances (PFASs)—A Synthesis of Current Knowledge with Proposal of Molecular Mechanism. IJMS 22, 2148 (2021).

46. Choi, S., Cho, N. & Kim, K. K. The implications of alternative pre-mRNA splicing in cell signal transduction. Exp Mol Med 55, 755–766 (2023).

47. Baralle, F. E. & Giudice, J. Alternative splicing as a regulator of development and tissue identity. Nat Rev Mol Cell Biol 18, 437–451 (2017).

48. Langfelder, P., Mischel, P. S. & Horvath, S. When Is Hub Gene Selection Better than Standard Meta-Analysis? PLoS ONE 8, e61505 (2013).

49. Horvath, S. & Dong, J. Geometric Interpretation of Gene Coexpression Network Analysis. PLoS Comput Biol 4, e1000117 (2008).

50. Ackerman, W. E. et al. Transcriptomics-Based Subphenotyping of the Human Placenta Enabled by Weighted Correlation Network Analysis in Early-Onset Preeclampsia With and Without Fetal Growth Restriction. Hypertension 80, 1363–1374 (2023).

51. Gan, L., Cookson, M. R., Petrucelli, L. & La Spada, A. R. Converging pathways in neurodegeneration, from genetics to mechanisms. Nat Neurosci 21, 1300–1309 (2018).

52. Lien, Y.-C. et al. Human Placental Transcriptome Reveals Critical Alterations in Inflammation and Energy Metabolism with Fetal Sex Differences in Spontaneous Preterm Birth. IJMS 22, 7899 (2021).

53. Isaac, R. et al. TM7SF3, a novel p53-regulated homeostatic factor, attenuates cellular stress and the subsequent induction of the unfolded protein response. Cell Death Differ 24, 132–143 (2017).

54. Isaac, R. et al. A seven-transmembrane protein-TM7SF3, resides in nuclear speckles and regulates alternative splicing. iScience 25, 105270 (2022).

55. Burton, G. J., Yung, H.-W., Cindrova-Davies, T. & Charnock-Jones, D. S. Placental Endoplasmic Reticulum Stress and Oxidative Stress in the Pathophysiology of Unexplained Intrauterine Growth Restriction and Early Onset Preeclampsia. Placenta 30, 43–48 (2009).

56. Capatina, N., Hemberger, M., Burton, G. J., Watson, E. D. & Yung, H. W. Excessive endoplasmic reticulum stress drives aberrant mouse trophoblast differentiation and placental development leading to pregnancy loss. The Journal of Physiology 599, 4153–4181 (2021).

57. Bindesbøll, C. et al. NBEAL1 controls SREBP2 processing and cholesterol metabolism and is a susceptibility locus for coronary artery disease. Sci Rep 10, 4528 (2020).

58. Burke, K. T. et al. Transport of maternal cholesterol to the fetus is affected by maternal plasma cholesterol concentrations in the Golden Syrian hamster. Journal of Lipid Research 50, 1146–1155 (2009).

59. Haug, M., Dunder, L., Lind, P. M., Lind, L. & Salihovic, S. Associations of perfluoroalkyl substances (PFAS) with lipid and lipoprotein profiles. J Expo Sci Environ Epidemiol 33, 757–765 (2023).

60. Grundmann, U., et al. Characterization of cDNA encoding human placental anticoagulant protein (PP4): homology with the lipocortin family. Proc. Natl. Acad. Sci. U.S.A. 85, 3708–3712 (1988).

61. Rand, J. H. The annexinopathies: a new category of diseases. Biochimica et Biophysica Acta (BBA) - Molecular Cell Research 1498, 169–173 (2000).

62. Degrelle, S. A., Gerbaud, P., Leconte, L., Ferreira, F. & Pidoux, G. Annexin-A5 organized in 2D-network at the plasmalemma eases human trophoblast fusion. Sci Rep 7, 42173 (2017).

63. Bogdanova, N. et al. A common haplotype of the annexin A5 (ANXA5) gene promoter is associated with recurrent pregnancy loss. Human Molecular Genetics 16, 573–578 (2007).

64. Sferruzzi-Perri, A. N., Sandovici, I., Constancia, M. & Fowden, A. L. Placental phenotype and the insulin-like growth factors: resource allocation to fetal growth. The Journal of Physiology 595, 5057–5093 (2017).

65. Shimada, H., Powell, T. L. & Jansson, T. Regulation of placental amino acid transport in health and disease. Acta Physiologica 240, e14157 (2024).

66. Fowden, A. L., Sferruzzi-Perri, A. N., Coan, P. M., Constancia, M. & Burton, G. J. Placental efficiency and adaptation: endocrine regulation. The Journal of Physiology 587, 3459–3472 (2009).

67. Hamburg-Shields, E. & Mesiano, S. The hormonal control of parturition. Physiological Reviews 104, 1121–1145 (2024).

68. Imai, K., Keele, L. & Tingley, D. A general approach to causal mediation analysis. Psychological Methods 15, 309–334 (2010).

69. Liu, J., Park, C., Li, K. & Tchetgen Tchetgen, E. J. Regression-based proximal causal inference. American Journal of Epidemiology 194, 2030–2036 (2025).

70. Richmond, R. C., Al-Amin, A., Davey Smith, G. & Relton, C. L. Approaches for drawing causal inferences from epidemiological birth cohorts: A review. Early Human Development 90, 769–780 (2014).

71. Tchetgen, E. J. T., Ying, A., Cui, Y., Shi, X. & Miao, W. An Introduction to Proximal Causal Learning. Preprint at 10.1101/2020.09.21.20198762 (2020).

72. Zivich, P. N. et al. INTRODUCING PROXIMAL CAUSAL INFERENCE FOR EPIDEMIOLOGISTS. American Journal of Epidemiology 192, 1224–1227 (2023).

73. Huang, J. Y. Leveraging multi-omic molecular negative controls for effect estimation in non-randomized human health and disease studies: a simulation study.

74. Tchetgen Tchetgen, E. The Control Outcome Calibration Approach for Causal Inference With Unobserved Confounding. American Journal of Epidemiology 179, 633–640 (2014).

75. Shi, X., Miao, W. & Tchetgen, E. T. A Selective Review of Negative Control Methods in Epidemiology. Curr Epidemiol Rep 7, 190–202 (2020).

76. Derisoud, E., Jiang, H., Zhao, A., Chavatte-Palmer, P. & Deng, Q. Revealing the molecular landscape of human placenta: a systematic review and meta-analysis of single-cell RNA sequencing studies. Human Reproduction Update 30, 410–441 (2024).

77. Ng, S. et al. Endocrine disrupting chemicals in maternal and umbilical cord plasma and their associations with birthweight in the GUSTO cohort. Environ Health 24, 57 (2025).

78. Framework for Estimating Noncancer Health Risks Associated with Mixtures of Per- and Polyfluoroalkyl Substances (PFAS).

79. Leader, J. et al. Transplacental transfer efficiency of environmental chemicals in association with gestational diabetes mellitus. Environment International 205, 109865 (2025).

80. Fitzgerald, E. et al. Hofbauer cell function in the term placenta associates with adult cardiovascular and depressive outcomes. Nat Commun 14, 7120 (2023).

81. Yong, H. E. J. et al. Increasing maternal age associates with lower placental CPT1B mRNA expression and acylcarnitines, particularly in overweight women. Front. Physiol. 14, 1166827 (2023).

82. Oxford Nanopore Technologies. Pychopper: cDNA read preprocessing and classification tool. (2024).

83. Mak, Q. X. C. Polishing De Novo Nanopore Assemblies of Bacteria and Eukaryotes With FMLRC2.

84. Li, H. New strategies to improve minimap2 alignment accuracy. Bioinformatics 37, 4572–4574 (2021).

85. Gao, Y. et al. ESPRESSO: Robust discovery and quantification of transcript isoforms from error-prone long-read RNA-seq data. Sci. Adv. 9, eabq5072 (2023).

86. Pardo-Palacios, F. J. et al. SQANTI3: curation of long-read transcriptomes for accurate identification of known and novel isoforms. Nat Methods 21, 793–797 (2024).

87. Chen, S., Zhou, Y., Chen, Y. & Gu, J. fastp: an ultra-fast all-in-one FASTQ preprocessor. Bioinformatics 34, i884–i890 (2018).

88. Patro, R., Duggal, G., Love, M. I., Irizarry, R. A. & Kingsford, C. Salmon provides fast and bias-aware quantification of transcript expression. Nat Methods 14, 417–419 (2017).

89. Jain, C., Koren, S., Dilthey, A., Phillippy, A. M. & Aluru, S. A fast adaptive algorithm for computing whole-genome homology maps. Bioinformatics 34, i748–i756 (2018).

90. Robinson, M. D., McCarthy, D. J. & Smyth, G. K. edgeR : a Bioconductor package for differential expression analysis of digital gene expression data. Bioinformatics 26, 139–140 (2010).

91. Baldoni, P. L. et al. Dividing out quantification uncertainty allows efficient assessment of differential transcript expression with edgeR. Nucleic Acids Research 52, e13–e13 (2024).

92. Langfelder, P. & Horvath, S. WGCNA: an R package for weighted correlation network analysis. BMC Bioinformatics 9, 559 (2008).

93. Love, M. I., Huber, W. & Anders, S. Moderated estimation of fold change and dispersion for RNA-seq data with DESeq2. Genome Biol 15, 550 (2014).

94. Risso, D., Ngai, J., Speed, T. P. & Dudoit, S. Normalization of RNA-seq data using factor analysis of control genes or samples. Nat Biotechnol 32, 896–902 (2014).

95. Ritchie, M. E. et al. limma powers differential expression analyses for RNA-sequencing and microarray studies. Nucleic Acids Research 43, e47–e47 (2015).

96. Antonov, M., et al. igraph enables fast and robust network analysis across programming languages. Preprint at 10.48550/ARXIV.2311.10260 (2023).

97. Yu, G., Wang, L.-G., Han, Y. & He, Q.-Y. clusterProfiler: an R Package for Comparing Biological Themes Among Gene Clusters. OMICS: A Journal of Integrative Biology 16, 284–287 (2012).

98. Carlson, M. org.Hs.eg.db. Bioconductor 10.18129/B9.BIOC.ORG.HS.EG.DB (2017).

