## Supplementary Figures 1-4 for "Natural variation in transplacental transfer efficiency exposes distinct transcriptional network architectures of PFAS effects on birth weight and gestational age"

**Contents**

**Supplementary Figure 1** – Concordance between PFBS-associated differential expression in placental explants and observational tissue samples

**Supplementary Figure 2** – PFAS effects on gestational age are mediated through co-expression network hubs rather than differentially expressed features.

**Supplementary Figure 3** – Maternal and fetal PFAS concentrations are correlated for low-TPTE but not high-TPTE compounds

**Supplementary Figure 4** – Network compartmentalization of PFAS-gestational age mediation effects scales with transplacental transfer efficiency (TPTE)

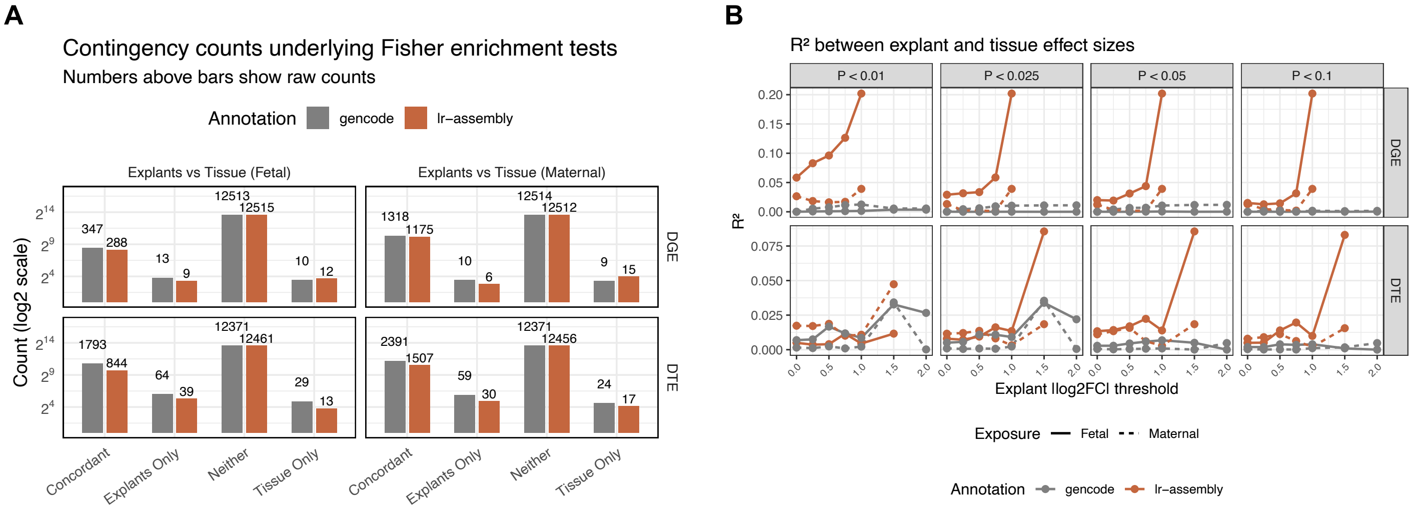


**Supplementary Figure 1 – Concordance between PFBS-associated differential expression in placental explants and observational tissue samples. (A)** Contingency counts from Fisher's exact tests of overlap between PFBS-associated features in placental explants (n = 18) and placental tissue (GUSTO, n = 124; n = 101 with fetal PFBS, n = 101 with maternal PFBS, n = 78 with paired measures), stratified by analysis level (rows: DGE = differential gene expression; DTE = differential transcript expression, where counts reflect genes with at least one differentially expressed isoform), exposure source (columns: fetal = cord blood PFBS; maternal = mid-gestation blood PFBS), and reference annotation (color: long-read assembly [terracotta]; GENCODE [grey]). Features are classified as concordantly significant in both datasets (Concordant), significant in explants only (Explants Only), significant in tissue only (Tissue Only), or in neither (Neither). The concordant overlap set was defined as nominally significant ($p < 0.05$) features with the same direction of effect in both datasets, and enrichment was tested against features nominally significant in explants ($|\log_2 FC| > 1$, $p < 0.05$) using Fisher's exact test. For isoform-level effects, overlap is assessed at the gene level: genes are counted if they harbor at least one significantly differentially expressed transcript in both datasets. Results are shown separately for fetal (cord blood) and maternal blood PFBS exposures. Models for placental tissue adjust for fetal sex and gestational age, while models for placental explants adjust for trimester (explants derived from $2^{nd}$ trimester and term samples). Both models adjust for technical variance estimated with RUVr. Raw counts are shown above bars; y-axis is on a $\log_2$ scale. **(B)** $R^2$ values quantifying correlation between PFBS effect sizes in explants and placental tissue are shown as a function of the $|\log_2 FC|$ threshold applied to explant differential expression results. Results are stratified by p-value threshold (columns), analysis level (rows: DGE = differential gene expression; DTE = differential transcript expression), reference annotation (color: long-read assembly [terracotta]; GENCODE [grey]), and exposure source (fetal = cord blood PFBS; maternal = mid-gestation blood PFBS). Only features passing the specified p-value and $|\log_2 FC|$ thresholds in explants with available measurements in GUSTO tissue samples were included in correlation calculations.

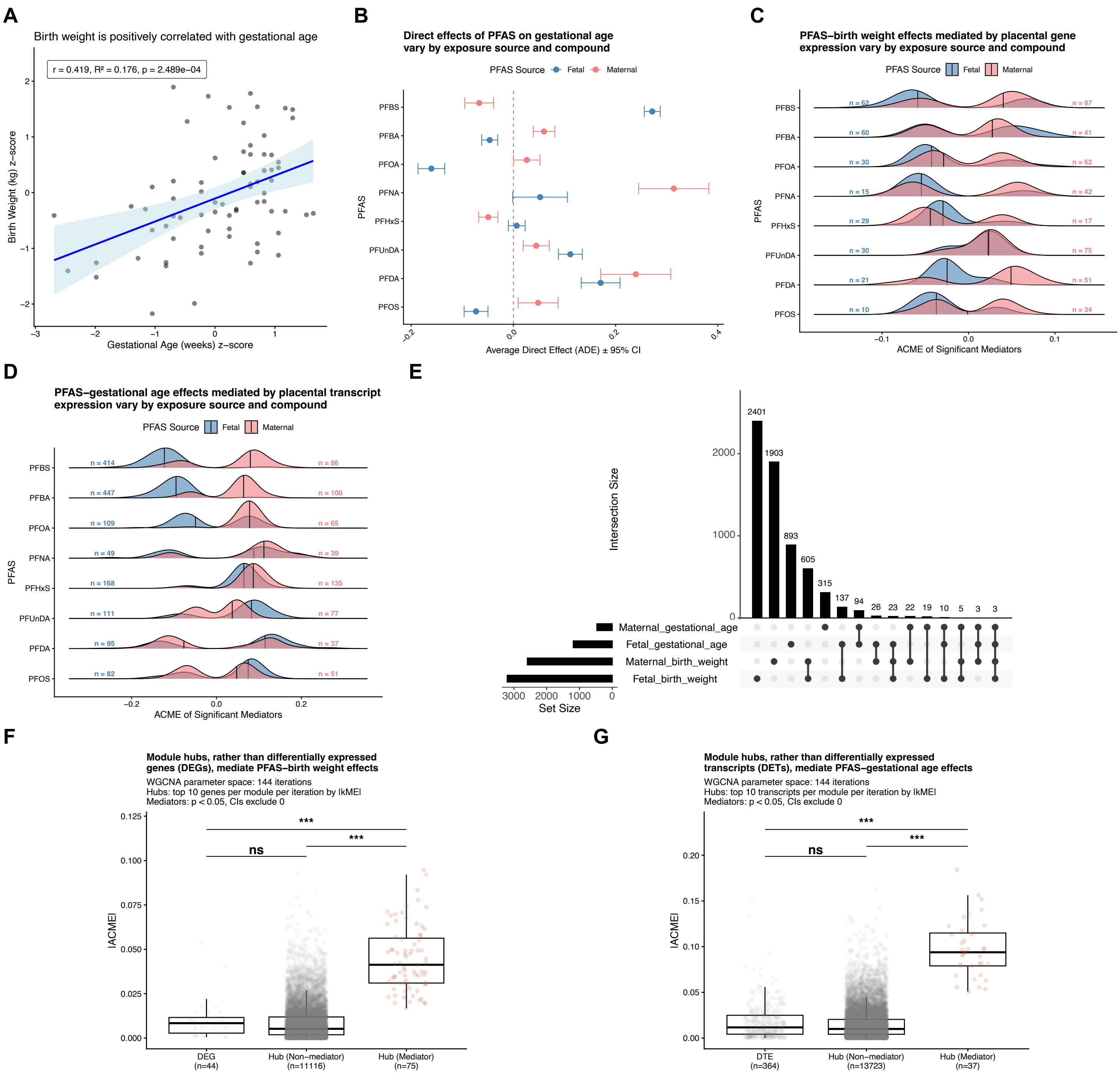


**Supplementary Figure 2 – PFAS effects on gestational age are mediated through co-expression network hubs rather than differentially expressed features.** **(A)** Pearson correlation between birth weight (kg) z-score and gestational age (weeks) z-score. **(B)** Average direct effects (ADE) of PFAS on gestational age vary by compound and exposure source. Mean ADE ± 95% confidence intervals for maternal (mid-gestation blood, pink points) and fetal (cord blood, blue points) PFAS exposure sources, calculated across all significant mediation pathways. Points represent weighted mean ADEs using inverse variance weighting across significant transcript mediators. PFAS compounds are ordered by transplacental transfer efficiency (TPTE). **(C)** PFAS–birth weight effects mediated by placental gene expression vary by exposure source and compound. Density distributions of absolute average causal mediation effects (|ACME|) for significant mediators ($p < 0.05$ and 95% CI excludes zero) by PFAS compound and exposure source. Blue distributions represent fetal (cord blood) PFAS exposures; pink distributions represent maternal (mid-gestation blood) PFAS exposures. Vertical lines within each distribution indicate the median |ACME|. Sample sizes indicate the number of significant mediating genes for each PFAS-source combination. **(D)** PFAS-gestational age effects mediated by placental transcript expression vary by exposure source and compound. Plot description as in **(C)**. **(E)** Overlap of significant mediator transcripts by exposure source and outcome. UpSet plot showing intersection sizes between sets of significant mediators across four conditions: fetal exposure-birth weight, maternal exposure-birth weight, fetal exposure-gestational age, and maternal exposure-gestational age. Horizontal bars (left) show total set sizes for each condition. Vertical bars (top) show the size of each intersection, with dots and connecting lines below indicating which sets are included in each intersection. Numbers above bars indicate the count of transcripts in each intersection. **(F)** Absolute average causal mediation effects (|ACME|) for birth weight mediator genes vs DEGs (FDR < 0.1, $|\log_2 FC| > 1$). Mediators stratified by network hubs (top 10 genes per module per iteration by |kME|). Statistical comparisons via Tukey HSD following ANOVA with bootstrapping (1000 iterations, n = 100 per category): ns = not significant, ***p < 0.001. **(G)** Absolute average causal mediation effects for gestational age mediator transcripts vs DETs. Plot description as in **(F)**.

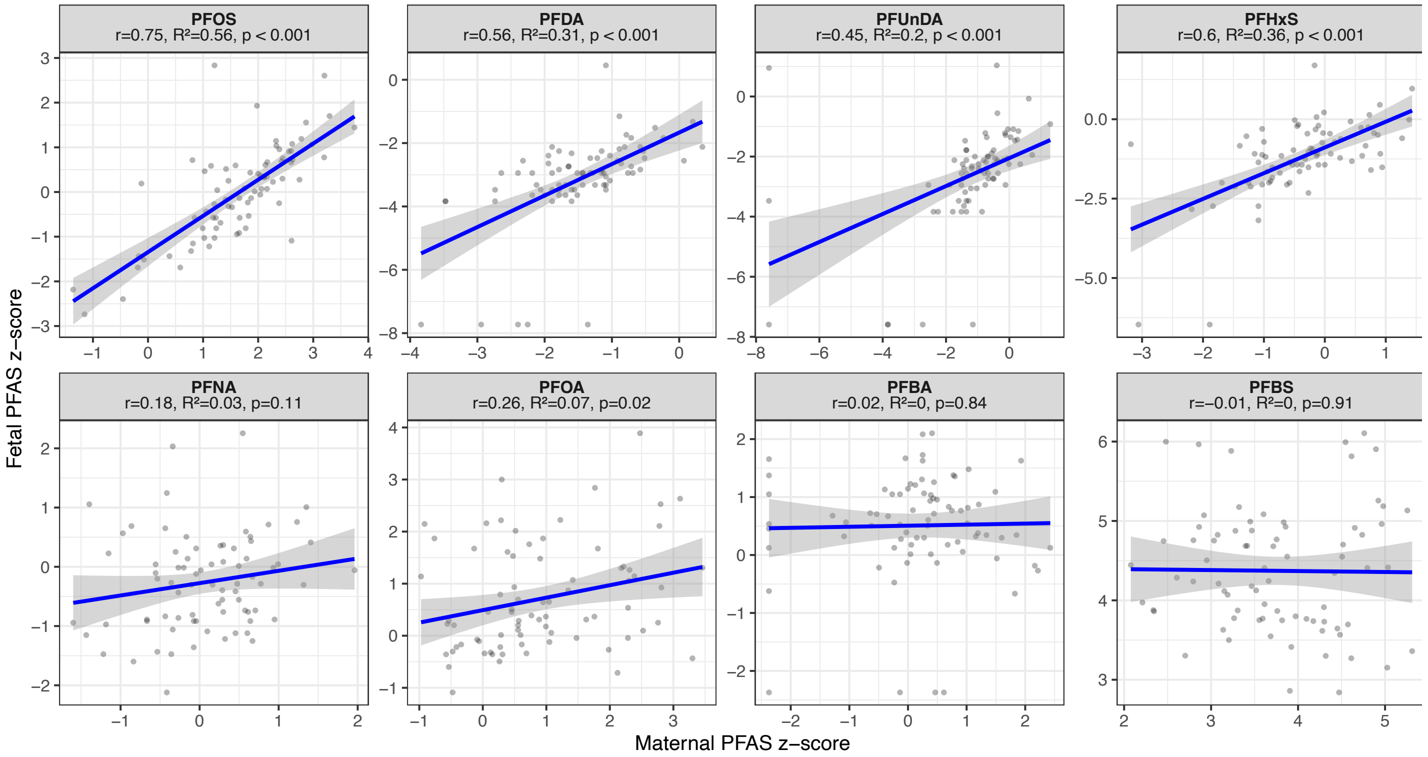


**Supplementary Figure 3 – Maternal and fetal PFAS concentrations are correlated for low-TPTE but not high-TPTE compounds.** Scatterplots of maternal blood (x-axis) and cord blood (y-axis) PFAS concentration z-scores for each of the eight measured compounds (n = 78 with paired measurements), ordered by transplacental transfer efficiency (TPTE). Pearson correlation coefficients, $R^2$, and p-values are shown for each compound.

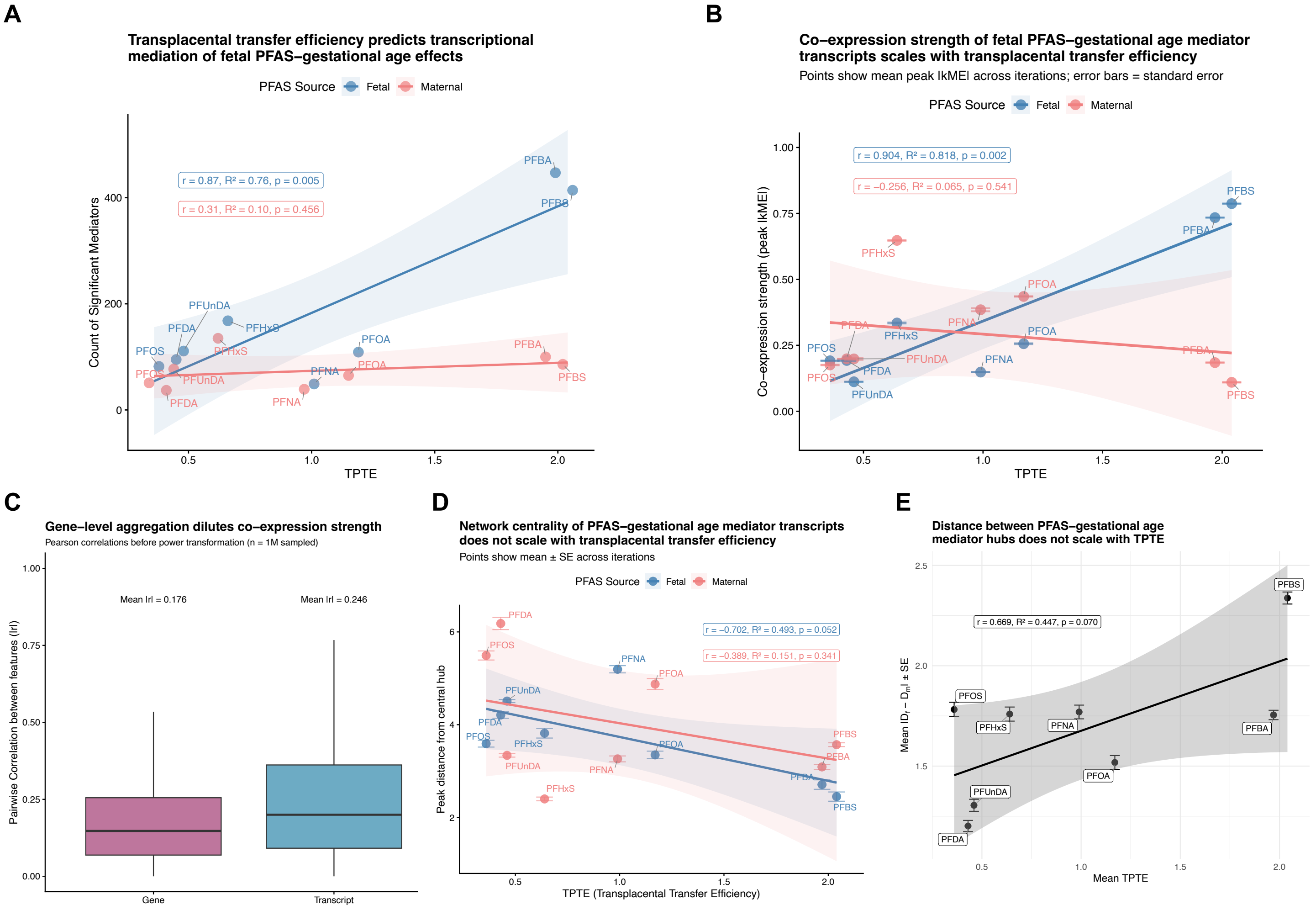


**Supplementary Figure 4 – Network compartmentalization of PFAS-gestational age mediation effects scales with transplacental transfer efficiency (TPTE).** Analyses of gestational age were restricted to spontaneous births (n = 72 with any PFAS measure; n = 42 with paired measures). **(A)** Count of significant mediators ($p < 0.05$, 95% CI excludes 0) across all modules and iterations plotted against TPTE for each PFAS compound and exposure source. Points represent total counts per PFAS-exposure combination. Blue points represent fetal (cord blood) exposures; pink points represent maternal (mid-gestation blood) exposures. Linear regression statistics shown separately for fetal and maternal exposures. **(B)** Peak absolute module eigengene correlation (|kME|) values (mean ± SE across iterations) plotted against TPTE for each PFAS compound and exposure source. Blue points represent fetal (cord blood) exposures; pink points represent maternal (mid-gestation blood) exposures. Error bars represent standard error. Linear regression statistics shown separately for fetal and maternal exposures. **(C)** Gene-level aggregation dilutes co-expression strength. Kernel density distributions of pairwise Pearson correlations (|r|) between features before power transformation, sampled from 1 million pairwise comparisons. Left panel shows gene-level correlations; right panel shows transcript-level correlations. **(D)** Distance from mediators to network hubs plotted against TPTE for each PFAS compound and exposure source. For each module-PFAS-exposure-iteration combination, peak distance is defined as the mode of the kernel density distribution of edge-weighted (1/r) shortest path lengths from the module's central hub transcript to all significant mediators, averaged across iterations. Blue points represent fetal (cord blood) exposures; pink points represent maternal (mid-gestation blood) exposures. Points show mean ± SE across iterations. Linear regression statistics shown separately for fetal and maternal exposures. **(E)** Absolute difference between fetal ($D_f$) and maternal ($D_m$) peak distances (mean ± SE across modules and iterations) plotted against TPTE for each PFAS compound. Points represent mean $|D_f - D_m|$ values; error bars represent standard error. Linear regression line and statistics shown.
